# Hepatitis E Virus-induced antiviral response by plasmacytoid dendritic cells is modulated by the ORF2 protein

**DOI:** 10.1101/2024.04.12.589258

**Authors:** Garima Joshi, Elodie Décembre, Jacques Brocard, Claire Montpellier, Martin Ferrié, Omran Allatif, Ann-Kathrin Mehnert, Johann Pons, Delphine Galiana, Viet Loan Dao Thi, Nolwenn Jouvenet, Laurence Cocquerel, Marlène Dreux

**Affiliations:** CIRI, INSERM, U1111, Université Claude Bernard Lyon 1, CNRS, UMR5308, École Normale Supérieure de Lyon, Univ Lyon, F-69007, Lyon, France; Université Claude Bernard Lyon 1, CNRS UAR3444, INSERM US8, ENS de Lyon, SFR Biosciences, Lyon 69007, France; Univ. Lille, CNRS, INSERM, CHU Lille, Institut Pasteur de Lille, U1019-UMR 9017-CIIL-Center for Infection and Immunity of Lille, F-59000 Lille, France; Institut Pasteur, Université de Paris, CNRS UMR 3569, Virus sensing and signaling Unit, 75015 Paris, France; Department of Infectious Diseases, Virology, Heidelberg University, Medical Faculty Heidelberg, Heidelberg, Germany and German Centre for Infection Research (DZIF), Partner Site Heidelberg, Heidelberg, Germany; Sup’biotech : École Des Ingénieurs En Biotechnologies, Villejuif, Paris

## Abstract

Type I and III interferons (IFN-I/III) are critical to protect the host during viral infection. Previous studies have shown that IFN-mediated antiviral responses against hepatitis E virus (HEV) are suppressed and defeated by viral escape mechanisms at play in infected hepatocytes. Here, we studied the anti-HEV function of IFN secreted by plasmacytoid dendritic cells (pDCs), which are specialized producers of IFNs. We showed that pDCs co-cultured with HEV-replicating cells secreted IFN in a cell-to-cell contact-dependent manner. Pharmacological inhibitor and antibodies targeting contact proteins revealed that pDC response against HEV required the endosomal nucleic-acid sensor TLR7 and adhesion molecules, such as ICAM-I and α_L_β_2_-integrin. IFNs secreted by pDCs reduced viral spread. Intriguingly, ORF2, the capsid protein of HEV, can be produced in various forms by the infected cells. During infection, a fraction of the intracellular ORF2 protein localizes into the nucleus while another ORF2 fraction packages viral genomes to produce infectious virions. In parallel, glycosylated forms of ORF2 are also massively secreted by infected cells. Using viral genome expressing ORF2 mutants, we showed that glycosylated ORF2 forms contribute to better recognition of infected cells by pDCs via regulation of contacts between infected cells and pDCs. ORF2 forms may thus modulate pDC-mediated anti-HEV response. Together, our results suggest that liver-resident pDCs, which exhibit comparable IFN-producing ability as blood-derived pDCs, may be essential to control HEV replication.

## Introduction

Hepatitis E virus (HEV) is the most common cause of acute viral hepatitis worldwide. It has been estimated that this virus infects approximately 100 million people every year and is responsible for 14 million symptomatic cases and 300,000 deaths, mainly in regions of the world with poor sanitary conditions **(Li et al., 2020)**. HEV infection is often asymptomatic and resolves on its own in case of healthy subjects. However, severe cases have mainly been reported in pregnant women, while chronic infections are more common in immunocompromised patients **(Lhomme et al., 2020)**. This makes host immunity a crucial factor in influencing the outcome of the disease. In addition, HEV infection is associated with a broad range of extrahepatic manifestations, including renal and neurological disorders **(Songtanin et al., 2023)**. The *Paslahepevirus balayani* species of the *Paslahepevirus* genus contains five HEV genotypes (gt) that are pathogenic in humans. HEV gt1 and gt2 are primarily transmitted through contaminated water, exclusively infect humans, and are responsible for waterborne hepatitis outbreaks in developing countries. In contrast, industrialized countries usually fall victim to HEV gt3 and gt4, which have a zoonotic origin **(Doceul et al., 2016)**. Chronic human infection with camel-associated HEV gt7 has also been reported **(Lee et al., 2016)**. Alarmingly, a rat HEV from the *Rocahepevirus* species, was recently reported to be also transmitted to humans **(Andonov et al., 2019; Sridhar et al., 2018)**.

The most common genotype causing chronic HEV infection in the developed world is gt3 **(Z. Ma et al., 2022)**. HEV infection has been recognized as a burgeoning issue in industrialized countries due to its chronicity in immunocompromised gt3-infected patients, the transmission of HEV through blood transfusion, a growing number of diagnosed HEV cases, and complications in patients with pre-existing liver disease **(Sayed et al., 2017)**. Importantly, HEV was recently ranked 6^th^ among the top 10 zoonotic viruses presenting the greatest risk of transmission to humans **(Grange et al., 2021)**.

HEV has a single-stranded, positive-sense RNA genome that contains three open reading frames (ORFs): ORF1, ORF2 and ORF3 **(Tam et al., 1991)**. ORF1 encodes the non-structural ORF1 polyprotein that displays domains essential for viral replication including the RNA-dependent RNA-polymerase (RdRp) **(Koonin et al., 1992)**. The RdRp produces a negative-sense RNA replicative intermediate, which serves as a template for the subgenomic RNA. The subgenomic RNA encodes both ORF2 and ORF3 proteins **(Graff et al., 2006)**. ORF2 protein is the viral capsid protein and ORF3 is a small multifunctional phosphoprotein, involved in particle egress **(Nimgaonkar et al., 2018)**.

ORF2, which is composed of 660 amino acids, is produced in 3 forms: infectious ORF2 (ORF2i), glycosylated ORF2 (ORF2g), and cleaved ORF2 (ORF2c) **(Montpellier et al., 2018)**. The precise sequences of ORF2i, ORF2g and ORF2c proteins have been proposed by two groups **(Ankavay et al., 2019; Hervouet et al., 2022; Montpellier et al., 2018; Yin et al., 2018)**. The ORF2i protein is not glycosylated and forms the structural component of infectious particles. In contrast, ORF2g and ORF2c proteins (herein referred to as ORF2g/c) are highly glycosylated, secreted in large amounts in the culture supernatant (*i.e.,* about 1000x more than ORF2i) and are the most abundant ORF2 forms detected in patient sera, where they are likely targeted by patient antibodies **(Ankavay et al., 2019; Montpellier et al., 2018)**. Thus, ORF2g/c forms may act as a humoral decoy that inhibits antibody-mediated neutralization due to their antigenic overlap with HEV virions **(Yin et al., 2018)**. Whether ORF2g/c proteins play a specific role in the HEV life cycle, and how ORF2 forms regulate the innate immune responses need to be elucidated. ORF2 forms are produced from different pathways, including a major one in which the ORF2 proteins are directed to the secretion pathway, where they undergo maturation, glycosylation, and are subsequently released in significant quantities. A fraction of cytosolic ORF2i proteins is delivered to the virion assembly sites **(Bentaleb et al., 2022; Hervouet et al., 2022)**, while another fraction of ORF2i proteins translocates into the nucleus of infected cells, presumably to regulate host immune responses **(Ankavay et al., 2019; Hervouet et al., 2022; Lenggenhager et al., 2017)**.

Viral genomes can be recognized by cytosolic sensors *i.e.,* RIG-I, MDA5, LGP2 **(Rehwinkel & Gack, 2020)**, whose activation leads to the induction of type I and III interferons (IFN-I/III) pathways. HEV has developed many mechanisms to inhibit the IFN response *via* its three ORFs, described as follows. HEV gt1 ORF1 protein blocks RIG-I and Tank-binding kinase (TBK)-1 ubiquitination in hepatoma cells, thereby suppressing the pathway of IFN induction **(Nan et al., 2014)**. The amino-terminal region of HEV gt3 ORF1, harboring a putative methyltransferase (Met) and a papain-like cysteine protease (PCP) functional domain, inhibits IFN-stimulated response element (ISRE) promoter activation by inhibiting STAT1 nuclear translocation and phosphorylation **(Bagdassarian et al., 2018)**. Moreover, HEV gt1 and gt3 ORF2 protein antagonizes IFN induction in HEV-replicating hepatocytes by inhibiting phosphorylation of the transcriptional regulator IRF3 **(Lin et al., 2019)**. The ORF3 protein of the HEV gt3 can also suppress IFN response by blocking STAT1 phosphorylation **(Dong et al., 2012)**.

In human hepatoma cells and primary hepatocytes, HEV infection induces only IFN-III production **(Wu et al., 2018; Yin et al., 2018)**, which is comparatively less potent at lower doses and at early time points than IFN-I **(Lazear et al., 2019)**. However, elevated expression of IFN-stimulated genes (ISGs), which are effectors of IFN-I/III, was detected in the whole blood of HEV-infected patients **(Moal et al., 2013; Sayed et al., 2017; Yu et al., 2010)**, as well as in experimentally infected mice engrafted with human hepatocytes and chimpanzee **(Sayed et al., 2017; Yu et al., 2010)**. It must also be noted that the infectious HEV, with a complete replication cycle, has been shown to be more sensitive to IFNα treatment than the subgenomic replicon **(Zhou et al., 2016)**. Therefore, the host immune system can presumably mount an immune response to fight off HEV despite the attenuation of immune signaling within the host cells. Plasmacytoid dendritic cells (pDCs), which are key producers of IFN, could be instrumental in effectively counteracting the evasion strategies employed by HEV within the liver microenvironment. As pDCs are resistant to virtually all viruses **(Silvin et al., 2017),** these do not express viral proteins that may block IFN induction. pDCs are an immune cell type known to produce up to 1000-fold more IFNs than any other cell type **(Reizis, 2019)**. They are thus pivotal for host control of viral infections **(Venet et al., 2023; Yun et al., 2021; Reizis, 2019; Webster et al., 2016).** They also recruit NK cells at the site of viral replication, favor virus-specific T cell responses **(Cervantes-Barragan et al., 2012; Reizis, 2019; Swiecki et al., 2010; Webster et al., 2016, 2018)**, and secrete a large panel of pro-inflammatory cytokines, *e.g.,* tumor necrosis factor alpha (TNFα) and interleukin-6 (IL-6) **(Reizis, 2019)**. pDC stimulation is mediated by the recognition of viral nucleic acid by Toll-like receptors (TLR)-7 and −9, which localize in the endo-lysosomal compartment **(Reizis, 2019)**. In the case of RNA viruses, TLR7-ligand engagement results in the formation of a signal complex comprising IRAK1 (interleukin-1 receptor-associated kinase 1), IRAK4, and TRAF6 (tumor necrosis factor receptor–associated factor 6), in a MyD88-dependent (myeloid differentiation primary-response gene 88) manner. This further triggers the translocation of IRF7 (interferon-regulatory factor 7) into the nucleus, where it can induce the transcription of type I IFN genes. In addition, the NF-κB (nuclear factor-κB) and MAPKs (mitogen-activated protein kinases) mediated signaling are also activated **(Gilliet et al., 2008)**. It has been shown that liver pDCs retain the ability to produce copious amounts of IFNα **(Doyle et al., 2019)**. Whether liver-resident pDCs contribute to the control of HEV replication is unknown. Here, by using hepatocyte-derived cell lines selected for their immunocompetence, we investigated the ability of HEV-replicating cells to stimulate pDCs, and defined how ORF2 forms contribute to this process.

## Results

### Immunocompetence of HEV cellular models

The magnitude of the type I and III interferons (herein referred to as IFN-I/III) responses against viruses varies among cell types. We thus first assessed the immunological robustness of three cell lines, namely Huh-7.5, HepG2/C3A and PLC3 cells, which are permissive to HEV **(Montpellier et al., 2018; Schemmerer et al., 2016; Shukla et al., 2011)**. Huh 7.5 cells are a subclone of the hepatoma Huh-7 cells that express an inactivate version of RIG-I and thus display an increased permissiveness to viral infection **(Blight et al., 2002; Sumpter et al., 2005)**. HepG2/C3A cells resemble liver parenchymal cells and were selected from the primary hepatoblastoma derived cell line HepG2 for cell-contact inhibition, leading to a more hepatocyte-like phenotype compared to the parental line **(Knowles et al., 1980)**. PLC3 cells are a subclone of the PLC/PRF/5 hepatoma cell line broadly used to study HEV **(Montpellier et al., 2018)**. We measured mRNA levels of two interferon-stimulated genes (*MXA* and *ISG15*) and two cytokines (*TNF* and *IL6*) in the three cell lines upon stimulation with polyI:C (**Fig. 1a-c**), which is a synthetic analog of dsRNA that activates TLR-3, when added to the cell culture, and it activates the cytosolic sensors RIG-I and MDA5 when transfected into cells. *MXA* and *ISG15* are representatives of the IFN-I/III responses induced *via* IRF3-mediated signaling while *TNF* and *IL-6* are induced by NF-κB-signaling. All three cell types were deficient in TLR3 activity as none of them responded to treatment with polyI:C (**Fig. 1a-c**). The expression of *MXA*, *ISG15, TNF* and *IL-6* remained unaffected upon polyI:C transfection in Huh-7.5 cells **(Fig. 1a)**. In contrast, the expression of these four genes was upregulated upon activation of RIG-I and MDA5 in both HepG2/C3A and PLC3 cells **(Fig. 1b-c)**. Treatment with different IFN types and TNFα has been shown to inhibit HEV replication to a certain extent **(Murata et al., 2020; Todt et al., 2016; Wang et al., 2016; Zhou et al., 2016)**. Consequently, we determined whether the three cell lines responded to stimulation by three recombinant cytokines: IFN-β, IFN-λ3 *(i.e.,* representatives of Type I and III IFNs, respectively) or TNFα. We found that IFN-β treatment induced the expression of *MXA* and *ISG15* in all three cell lines but barely induced the proinflammatory cytokines, *TNF* and *IL-6* **(Fig. 1a-c)**. The three cell lines responded to the treatment with recombinant IFN-λ with an upregulation of *MXA* and *ISG15* mRNA (**Fig. 1a-c**). However, only HepG2/C3A cells showed significant induction of *IL-6* upon IFN-λ treatment **(Fig. 1b)**. Treatment with TNFα mainly upregulated the proinflammatory cytokine *TNF* in the three cell lines **(Fig. 1a-c)**. Together, our results show that HepG2/C3A and PLC3 cells are sufficiently immunocompetent and were therefore selected for further investigation.

**Figure 1.**
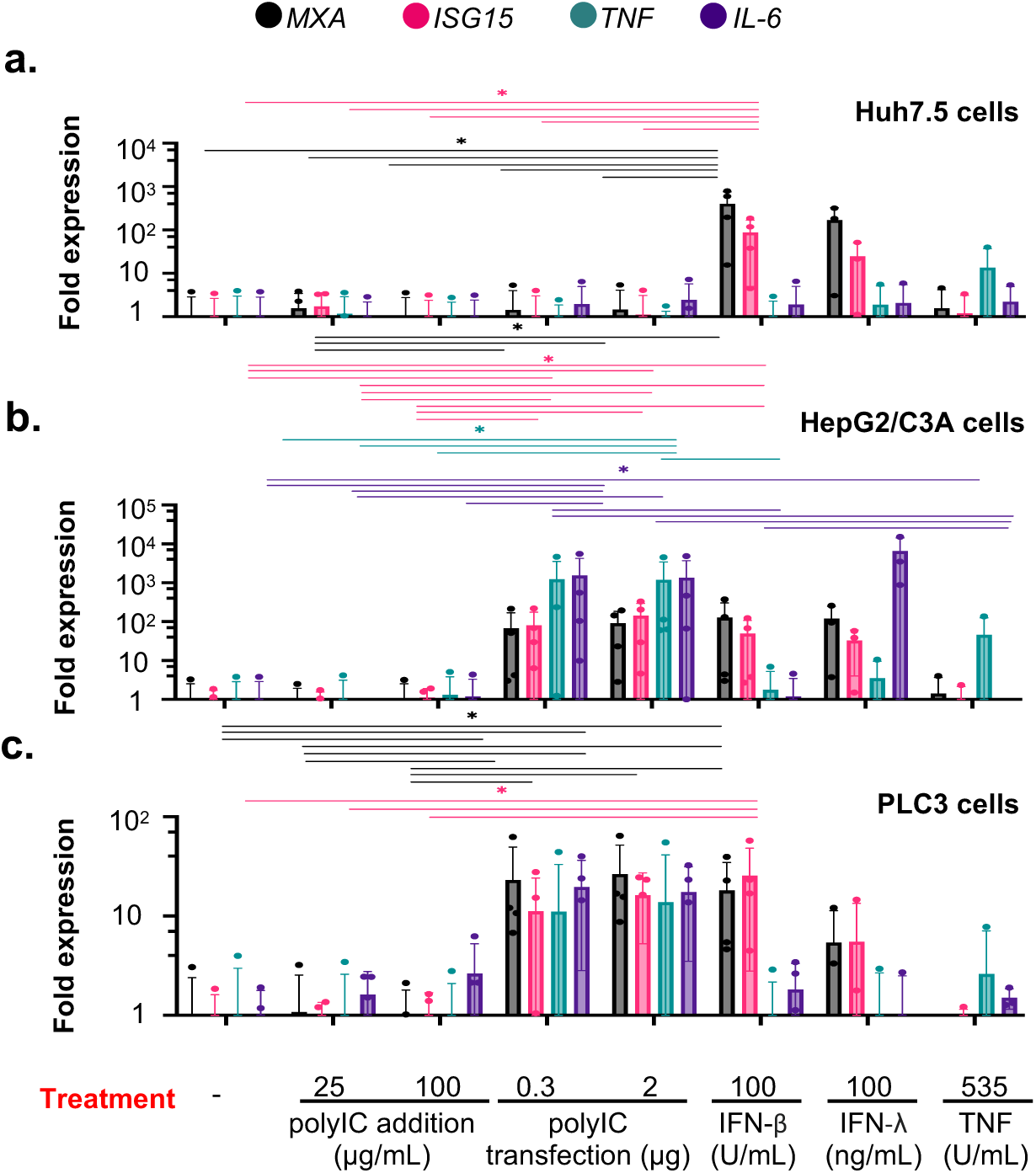
Immuno-responsiveness of cell types selected as highly susceptible to HEV infection. **(A-C)** Quantification of transcript expression levels of representatives of IFN-I/λ-related pathway, *i.e., MxA and ISG15*, NF-KB-induced signaling, *i.e., TNF* and *IL-6*, upon stimulation by agonists of TLR-3 *i.e.,* addition of poly(I:C); RIG-I/MDA-5 cytosolic sensors by transfection of LMW poly(I:C) and addition of recombinant IFN-β, IFN-λ3, TNF at the indicated concentrations for 6 hour-incubation, determined by RT-qPCR. Analyses were performed in Huh-7.5 cells (**A**), HepG2/C3A cells (**B**) and PLC3 cells (**C**); bars represent mRNA copy number per µg total RNA; means ± standard deviation (SD); each dot represents one independent experiment, *n*[=[4 for treatments including polyI:C addition/transfection and IFN-β treatment, n=3 for treatment with IFN-λ and TNF; statistical analysis using paired pairwise wilcox test; *p* values as: * ≤0.05, ** ≤0.005 and *** ≤0.0005.

Next, PLC3 and HepG2/C3A cells were electroporated with capped RNA of the HEV gt3 p6 strain **(Shukla et al., 2012)**. RT-qPCR analyses showed that HepG2/C3A and PLC3 cells produced a significant amount of viral RNA **(Supplementary Fig. 1a-b)** compared to mock-electroporated cells (Control). Immunostaining of ORF2 protein in PLC3 and HepG2/C3A cells at 6 days post-electroporation (d.p.e.) confirmed viral replication **(Supplementary Fig. 1c)**. In addition, we optimized a staining protocol to quantify ORF2-positive cells by flow cytometry. Flow cytometry analyses showed that about 50% of PLC3 cells were positive for ORF2 at 6 d.p.e. **(Supplementary Fig. 1d)**. Overall, these results indicated that HepG2/C3A and PLC3 cells are thus suitable models to perform immunological studies in the context of HEV replication.

### Activation of pDCs upon contact with HEV-replicating cells leads to antiviral response

Next, we looked at expression levels of two ISGs and three cytokines in HEV-replicating HepG2/C3A and PLC3 cells (**Fig. 2a-b**). We found a significant increase in mRNA levels of 3 ISGs (*ISG15*, *IL-6*, *IFN-λ1*) in HepG2/C3A cells at 6 d.p.e. **(Fig. 2a**; left panel). *OAS2,* which is known to activate RNase L antiviral activity, was also induced in HepG2/C3A (albeit not significantly). In PLC3 cells, mRNA abundance of *MXA*, *ISG15*, *IL-6, OAS2* and *IFN-λ1* remained unchanged **(Fig. 2b**; left panel). These results are in accordance with the higher responsiveness of HepG2/C3A cells to activation of cytosolic sensors and cytokines as compared to PLC3 cells **(Fig. 1b-c)**. Notably, the level of *IFN-λ1* mRNA, which has known anti-viral role against HEV in mice **(Sari et al., 2021)**, increased upon viral replication only in HepG2/C3A cells **(Fig. 2a-b**; left panels). In addition, *TNF* expression was slightly inhibited in HEV-replicating PLC3 cells **(Fig. 2b**; left panel) in accordance with previous results **(Hervouet et al., 2022).** These results showed that, although HepG2/C3A and PLC3 cells are immunocompetent upon stimulation (**Fig. 1**), IFN-I/III response is only modestly induced upon HEV replication in HepG2/C3A cells and marginally in PLC3 cells. In line with previous *in vitro* studies **(Bagdassarian et al., 2018; Dong et al., 2012; Lin et al., 2019; Nan et al., 2014)**, these results suggest that HEV is a poor inducer of IFN response.

**Figure 2.**
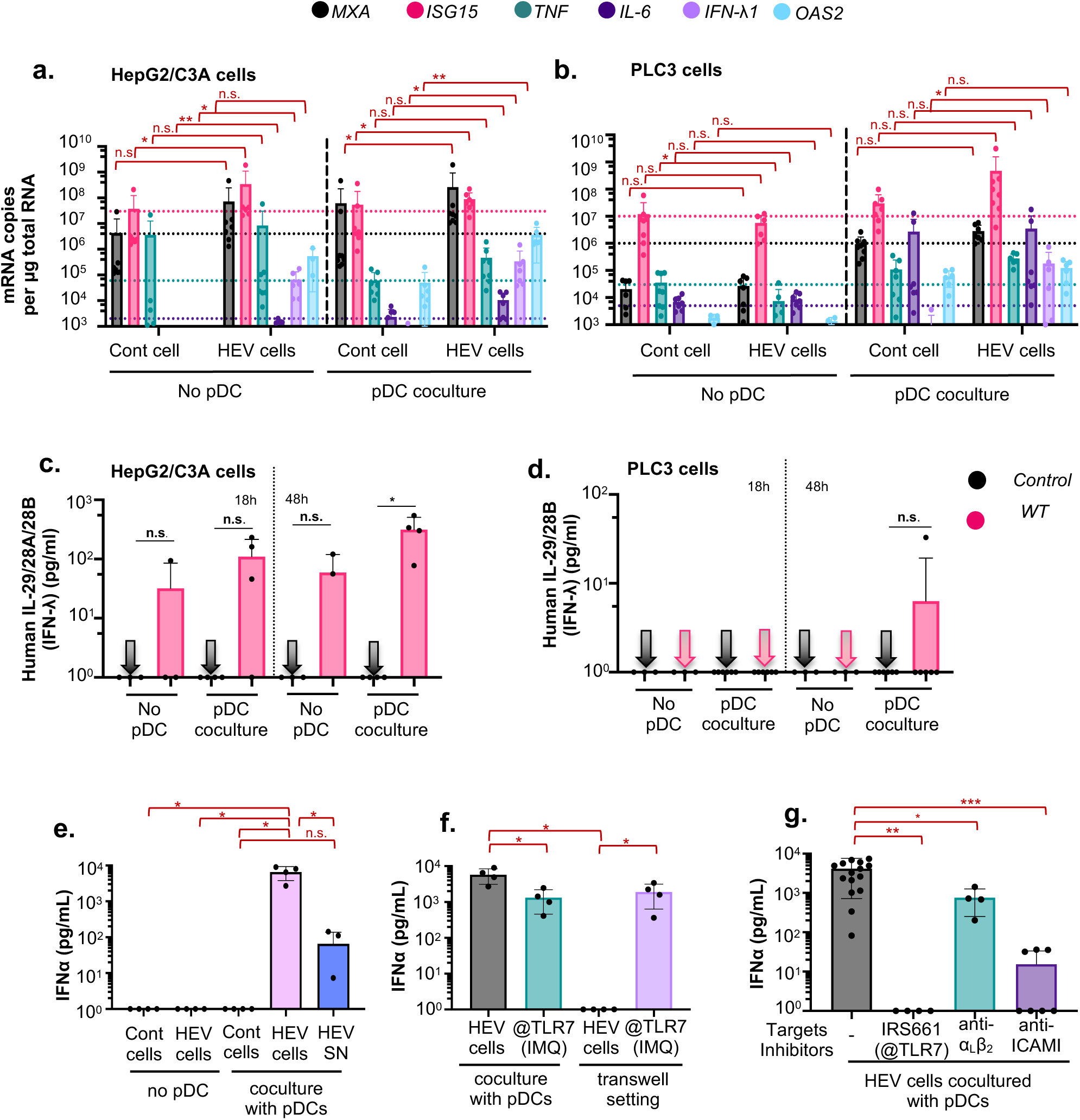
pDC response against HEV infected cells. (**A-B**) Quantification of the transcript expression levels of representatives of IFN-I/λ-signaling (*i.e., MxA, ISG15,OAS2 and IFNλ1*) and NF-KB-related pathway (*i.e., TNF and IL-6*) determined at 6 d.p.e. of HepG2/C3A (**A**) and PLC3 **(B)** cells that were electroporated with 10µg RNA [HEV cells] or mock electroporated without RNA [Cont cells], in the absence (left panels) or in co-culture with pDCs for 18 hours (right panels), as indicated. No pDC and pDC co-culture conditions have been separated into left and right panels as steady state levels of gene expression are different in these two datasets and therefore cross-comparisons between the two panels must be avoided. Bars represent copy number per µg total RNA; means ± SD; each dot represents one independent experiment (n=3 to 8). Statistical analysis was done using Wilcoxon rank sum test with continuity correction; *p* values as: * ≤0.05, ** ≤0.005 and *** ≤0.0005. **(C-D)** pDCs were cocultured with HEV-electroporated PLC3 or HepG2/C3A cells for 18 hours and 48 hours. Quantification of IL-29/28A/28B in supernatants of pDCs co-cultured with HEV-replicating HepG2/C3A **(C)** and PLC3 cells **(D)**; n≥3; Statistical analysis was done using Wilcoxon rank sum test with continuity correction; *p* values as: * ≤0.05, ** ≤0.005 and *** ≤0.0005. **(E-G)** pDCs were co-cultured with HEV-electroporated PLC3 cells, and their supernatants, in various settings or treated by inhibitors, as indicated, for 18 hours. **(E)** Quantification of IFNα in supernatants of pDCs co-cultured with HEV-replicating PLC3 cells [HEV cells] or treated with supernatants from HEV infected cells [HEV SN] *versus* in the absence of pDCs [no pDC]; the uninfected [cont] cells were electroporated without HEV RNA and used as negative control. Bars represent means ± SD and each dot represents one independent experiment (n=4). (**F**) Quantification of IFNα in supernatants of pDCs in co-culture or separated from HEV-electroporated PLC3 cells by a permeable membrane (0.4 μm) of transwell [transwell setting]. The TLR7 agonist, imiquimod [IMQ] was used as positive control. Results presented as in **E**; (n=4). (**G**) Co-culture of pDCs and HEV-electroporated PLC3 cells were treated by inhibitors of TLR7 [IRS661], blocking antibodies against α_L_β_2_-integrin and ICAM-1, followed by the quantification of IFNα in supernatants of the co-cultures. Results presented as in **E**; n>3 independent experiments. Statistical analysis was done using Wilcoxon rank sum test with continuity correction; *p* values as: * ≤0.05, ** ≤0.005 and *** ≤0.0005

*In vivo* studies of HEV infection (i.e., at tissue level) have, nonetheless, demonstrated induction of antiviral responses **(Moal et al., 2013; Sayed et al., 2017; Yu et al., 2010)**. We thus thought to investigate whether human primary pDCs can mount the IFN-I/III response. The human primary pDCs were stimulated upon co-culture for 18 hours with HEV-replicating cells. As in aforementioned experiments, we analyzed mRNA abundance of *MXA, ISG15, IFN-* λ*1*, *OAS2 TNF, and IL-6* by RT-qPCR **(Fig. 2a-b**; right panels). We observed upregulated expression of all these effectors, and among these, *MXA*, *ISG15, IFN-* λ*1* and *OAS2* were significantly upregulated when pDCs were co-cultured with HEV-replicating HepG2/C3A cells, as compared to basal levels with uninfected cells (**Fig. 2a-b, right panels**). Overall, likely due to the inhibitory mechanisms of HEV viral products in PLC3 cells targeting IFN-I/III amplification pathways, there was a poor activation of downstream effectors, ISGs and cytokines in the mixed cell culture, with the exception of *IFN-* λ*1*. The induction of *IFN-* λ*1* was common to the pDC co-cultures of HEV-replicating HepG2/C3A and PLC3 cells. We tested if this was true at the protein level by ELISA for lambda IFNs or type III IFNs (IFN-λ1[IL-29], IFN-λ2 [IL-28A] and IFN-λ3 [IL-28B]). We observed that IFN-III protein levels were significantly upregulated only at 48h post pDC co-culture with HEV-replicating HepG2/C3A cells, as compared to control cells co-cultured with pDCs **(Figure 2c)**. Even though IFN-λ1 mRNA levels were upregulated upon co-culture of pDCs with HEV-replicating PLC3 cells as early as 18 hours of coculture **(Figure 2b; right panel)**; only a small amount of secreted IFN-III was detectable at the protein level 48 hours post co-culture **(Figure 2d)**. This is possibly because of the lesser sensitivity of the ELISA over RT-qPCR analyses (i.e., lower dynamic range). Further analyses with more sensitive/advanced approaches for IFN-III detection will be needed. Therefore, IFN-III is more robustly upregulated in pDC coculture of HEV-replicating HepG2/C3A cells.

Next, we tested whether pDCs can produce IFNs in response to HEV-replicating cells by quantifying the secreted levels of multiple subtypes of IFNα as representative of IFN-I signaling **(Fig. 2e)**. HEV-replicating PLC3 cells alone did not produce detectable IFNα levels, in accordance with previous findings showing that HEV replication does not induce IFN-I secretion **(Moal et al., 2013; Nan et al., 2014; Sayed et al., 2017; Yu et al., 2010)**. In contrast, when pDCs were co-cultured with HEV-replicating PLC3 cells, more than 1000 pg/ml of IFNα were secreted, whereas no IFNα was detected when pDCs were co-cultured with uninfected control cells **(Fig. 2e)**. We also treated pDCs with cell-free HEV to verify if circulating virus particles can also activate pDCs in a cell-independent manner. pDCs exposed to the filtered supernatant [SN] collected from HEV-replicating cells very modestly secreted IFNα (Fig. 2e), suggesting that physical contact between pDCs and HEV-replicating cells is required for the robust pDC IFN-I secretion. Therefore, we assessed whether HEV-replicating PLC3 cells activated pDCs when the two cell types were separated by a 0.4 μm-permeable membrane, which allows diffusion of virions but not cells. In this Transwell setting, pDCs placed in the top chamber did not produce any IFNα, validating that physical contact between the cells is required for pDC stimulation **(Fig. 2f)**. Imiquimod [IMQ], which is a soluble agonist of TLR7 **(Gibson et al., 2002)**, was used as an additional positive control to rule out non-specific effect of the setting **(Fig. 2f)**. Furthermore, when the co-cultures were treated with blocking antibodies against ICAM-I and α_L_β_2_-integrin, two cell-cell adhesion proteins that mediate cellular contacts **(Assil, Coléon, et al., 2019; Marlin & Springer, 1987)**, pDC response to infected cells was significantly reduced as compared to untreated pDCs in co-culture with HEV-replicating cells **(Fig. 2g)**. Since TLR7 is responsible for sensing replicative viral RNAs in pDCs **(Assil, Coléon, et al., 2019; Reizis, 2019; Webster et al., 2016)**, we tested its contribution in pDC response to HEV-replicating cells. When the co-cultures were treated with the TLR7 inhibitor IRS661, no IFNα was secreted (**Fig. 2g**), suggesting that stimulation of pDCs by HEV-replicating cells is mainly mediated by TLR7. Together, our results show that direct contacts mediated by ICAM-I and, at least in part, with α_L_β_2_-integrin, enable pDCs to sense and respond to HEV-replicating cells in a TLR7-dependent manner.

We then tested whether IFNα produced by pDCs inhibited viral propagation in either HepG2/C3A or PLC3 cells **(Fig. 3a-c)**. Upon 18 hours of co-culture of pDCs with HEV-replicating HepG2/C3A cells, we observed a 1-log decrease in viral RNA yield (readout for viral replication) compared with the absence of pDCs **(Fig. 3a)**. At this early time of co-culture, the impact of pDCs on viral RNA replication was not yet observed for PLC3 cells **(Fig. 3b)**. This may result from a stronger upregulation of ISGs in HEV-replicating HepG2/C3A cells (in absence of pDC) compared to PLC3 cells, which may result in the synergy of the antiviral impact of pDCs, leading to earlier decrease in viral replication in HepG2/C3A cells. As we did not observe pDC-mediated viral control in HEV-replicating PLC3 cells at this early time point, we thus studied the effect of pDCs when co-cultured for 48 hours. After a longer co-culture of 48 hours with pDCs, we observed a 50% reduction of HEV-replicating cells as compared to cultures without pDCs **(Fig. 3c).**

**Figure 3.**
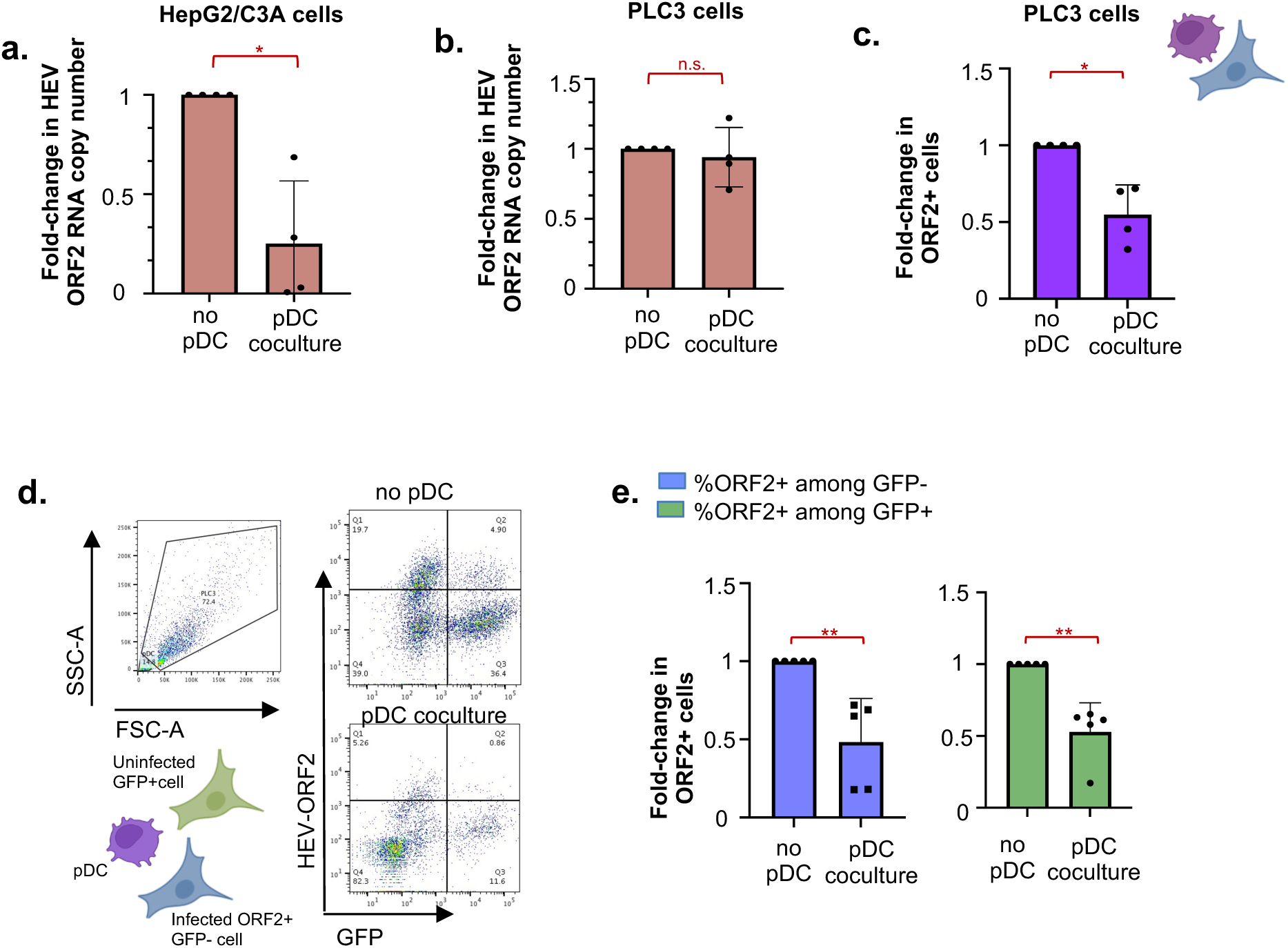
Control of HEV infection by pDCs. **(A-B)** Quantification of HEV RNA replication levels in HepG2/C3A (**A**) and PLC3(**B)** cells, in the absence or in co-culture with pDCs, as indicated; means ± SD; each dot represents one independent experiment in terms of fold expression compared to [no pDC] condition; (n=4) for HepG2/C3A cells and PLC3 cells. Statistical analysis was done using Wilcoxon rank sum test with continuity correction and p value adjustment with Bonferroni method; p values as: * ≤0.05, ** ≤0.005 and *** ≤0.0005. **(C)** Quantification of ORF2 expressing cells in the presence or absence of pDCs by flow cytometry; Fold-change in HEV-replicating (ORF2+) PLC3 cells in the absence and presence of pDCs for 48 hours (n=4). **(D-E)** HEV-replicating PLC3 cells (GFP^-^ ORF2^+^) and uninfected cells (GFP^+^ ORF2^-^) were co-cultured in the presence and absence of pDCs for 48 hours. Quantification of ORF2 and GFP expressing cells by flow cytometry; **(D)** Results are presented as representative dot blots. (**E**) Fold-change in ORF2^+^ GFP^-^ and ORF2^+^ GFP^+^ cells was quantified by flow cytometry. Bars represent means ± SD and each dot per independent experiment (n=5). Statistical analysis was done using Wilcoxon rank sum test with continuity correction and p value adjustment with Bonferroni method; p values as: * ≤0.05, ** ≤0.005 and *** ≤0.0005.

To further examine whether the pDC-mediated IFN response decreases the percentage of newly infected cells, we co-cultured a mixture of PLC3 cells, electroporated with HEV RNA and not expressing GFP (ORF2+/GFP- cells) along with uninfected cells positive for GFP (ORF2-/GFP+) in the presence or absence of pDCs for 48 hours. The impact of pDC response on HEV spread was assessed by flow cytometry to quantify the percentage of newly infected cells, i.e., GFP+ cells that became ORF2+ due to viral spread (ORF2+/GFP+) in the presence *versus* absence of pDCs **(Fig. 3d-e)**. Infected cells were distinguished from pDCs on the basis of size (FSC-SSC gating), and then the infected cell type (PLC3 cells) was gated for expression of ORF2 and/or GFP **(Fig. 3d)**. The results demonstrated that pDC response reduced viral replication in cells replicating HEV **(Fig. 3e**, blue bars) as well as controlled viral spread to naive GFP^+^ cells by half, upon prolonged co-culture **(Fig. 3e**, green bars). Collectively, our results showed an effective control of HEV spread by pDC-mediated antiviral activities involving IFN-I production.

### Modulation of pDC response by HEV expressing ORF2 mutant is dependent on the host cell type

pDCs responded to contact with HEV-replicating cells *via* TLR7 recognition (**Fig. 2e**), suggesting that HEV genome is transferred from infected cells to the endosomal compartments of pDCs, where TLR7 localizes **(Dreux et al., 2012)**. The distinctive feature of the HEV ORF2 capsid protein is its generation in three distinct forms (ORF2i, ORF2g, and ORF2c), each with different localization and trafficking patterns within infected cells **(Ankavay et al., 2019; Hervouet et al., 2022; Lenggenhager et al., 2017)**. To address whether these distinct ORF2 forms impact the host immune responses in infected cells, and consequently and/or additionally the sensing of infected cells by pDCs, we tested a series of ORF2 protein mutants of the p6 HEV strain (WT) **(Supplementary Fig. 1c)**. The first mutant, called 5R/5A mutant, expresses an ORF2 in which the arginine-rich motif (ARM) located in the ORF2 N-terminal region, serving as a nuclear localization signal, was mutated and thus prevents ORF2 translocation into the nucleus **(Hervouet et al., 2022)**. The NES mutant expresses an ORF2 that lacks one of its nuclear export signals (NES), leading to its retention in the nucleus **(Hervouet et al., 2022)**. Glycosylated proteins can influence immunological functions *via* their recognition by C-type Lectin receptor (CLRs) **(Cambi et al., 2005)**, either by weakening TLR7 and TLR9-mediated IFN-I/III response **(Florentin et al., 2012; Meyer-Wentrup et al., 2008)**, or contributing to cell interaction and viral uptake **(Bermejo-Jambrina et al., 2018)**. Therefore, we generated an additional mutant, called STOP mutant, expressing an ORF2 protein with a non-functional signal peptide, preventing its translocation into the endoplasmic reticulum and thus production of glycosylated ORF2g/c forms, as recently described **(Nagashima et al., 2023)**. This mutant carries a stop codon in the ORF2 signal peptide, which does not affect ORF3 expression. In the STOP mutant, ORF2 protein translation starts at the first initiation codon (Met^1^), stops at stop codon (*^10^) and restarts at the second initiation codon (Met^16^) (**Supplementary Fig. 2a**). PLC3 cells expressing this mutant produce intracellular ORF2i form (**Supplementary Fig. 2b**, right panel), efficiently replicate HEV genome, (**Supplementary Fig. 2d**) and produce HEV particles (**Supplementary Fig. 2c**, IP P1H1 and **Supplementary Fig. 2e-f**), but no ORF2g/c proteins (**Supplementary Fig. 2c**, IP P3H2). It is noteworthy that we observed the absence of ORF2g/c production in cell culture supernatants of the STOP mutant expressed in both PLC3 and HepG2/C3A cells **(Supplementary Figure 2g)**. Finally, a previously characterized replication-defective p6 mutant (GAD) was also included in the analysis **(Emerson et al., 2013)**.

The subcellular distribution of ORF2 mutants, compared with wild-type (WT) ORF2, was first characterized using confocal microscopic analyses. The WT ORF2 was found in both cytoplasm and nucleus, in both HepG2/C3A and PLC3 cells **(Fig. 4a-b).** The 5R/5A mutant was excluded from the nucleus whereas the NES mutant accumulated in the nucleus in both cell types (**Fig. 4a-b**), in accordance with previous observations **(Ankavay et al., 2019; Hervouet et al., 2022)**. The STOP mutant exhibited a localization similar to that of WT ORF2 (**Fig. 4a-b**). As expected, the GAD mutant, in which the polymerase active site was mutated and thus is replication-defective, did not produce ORF2 (**Fig. 4a-b**). We then assessed viral RNA yield produced in cells electroporated with the four mutant genomes or with WT genome using RT-qPCR analysis. This attested comparable levels of HEV RNA, which is a prerequisite for testing the impact of this mutant panel on the response of co-cultured pDC. The HEV RNA levels were similar across the mutants and WT genomes at 6 d.p.e. in the HepG2/C3A cells **(Fig. 4c)**. PLC3 cells electroporated with the WT, 5R/5A, NES and STOP mutant genomes also yielded similar levels of viral RNAs at 6 d.p.e. (**Fig. 4d)**. About two-log less viral RNA was recovered in HepG2/C3A and PLC3 cells expressing the GAD genome mutant **(Fig. 4c-d).** Viral RNA detected in these cells likely represent ‘input’ RNA, i.e., electroporated viral RNA.

**Figure 4.**
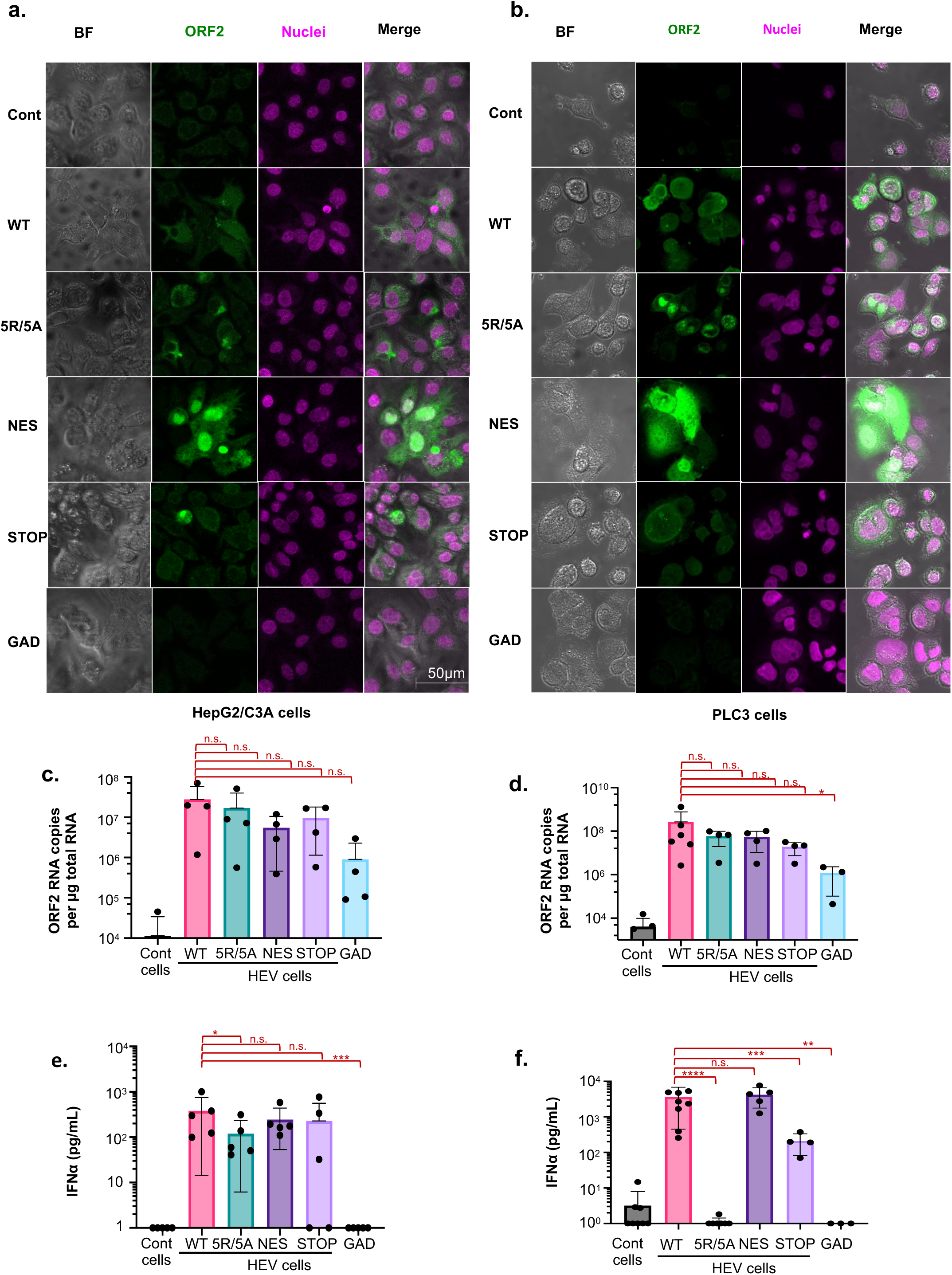
Impact of ORF2 nuclear localization on pDC activation by studying co-cultures with specific HEV mutants. Mutations were designed to inactivate: *i/* nuclear translocation of ORF2 as [5R5A] mutant, *ii/* the ORF2 export from nucleus as [NES] mutant and iii/ ORF2g/c secretion as [STOP] mutant. Cells harboring the [GAD] mutant that is deficient for viral replication and/or uninfected [cont] cells were used as negative controls. Plasmids containing the above-mentioned mutations in HEV genome *versus* WT, served as templates for *in vitro* transcription into HEV RNA, then transfected in either HepG2/C3A or PLC3 cells, as indicated. **(A-B)** Intracellular distribution of ORF2 (1E6 antibody, followed by targeting secondary antibody with AlexaFluor488) in HepG2/C3A (**A**) or PLC3 **(B)** cells electroporated with different mutants of HEV. (**C-D**) Quantification of HEV replication levels of mutants *versus* WT in HepG2/C3A (n=4) (**C**) or PLC3 cells (n=3 to 6) (**D**). Results represent HEV RNA copy number (primer/amplicon designed in *ORF2* RNA) per µg total RNA, detected by RT-qPCR; bars represent means ± SD and dot represent independent experiments. (**E-F**) Quantification of IFNα in supernatants of pDCs that were co-cultured with cells harboring HEV with the indicated mutant genome *versus* WT, and GAD or uninfected, as negative controls; for 18 hours; 6 d.p.e. of HepG2/C3A cells (n=5) (**E**) and PLC3 cells (n=3 to 8) (**F**); bars represent means ± SD and dot represent independent experiments. Statistical analysis was done using Wilcoxon rank sum test; *p* values as: * ≤0.05, ** ≤0.005 and *** ≤0.0005.

Quantification of IFNα production by pDCs co-cultured with HepG2/C3A expressing the different mutants demonstrated significant differences. First, pDCs co-cultured with HepG2/C3A cells expressing the GAD mutant did not trigger a pDC response **(Fig. 4e)**, nor in the context of PLC3 cells **(Fig. 4f)**. Since HEV RNA levels were reduced for GAD mutant, it is tempting to hypothesize that active viral replication is required for mounting a pDC-mediated viral response. Next, pDCs co-cultured with HepG2/C3A expressing 5R/5A demonstrated a significant reduction of IFNα production as compared to WT and 5R/5A *versus* NES **(Fig. 4e)**. These results suggested that, in this cellular model, the nuclear localization of ORF2 influences pDC stimulation. Along the same lines, pDC-mediated IFNα response was completely abolished upon co-culture with PLC3 cells expressing the 5R/5A mutant **(Fig. 4f)**. Since comparable HEV RNA levels were quantified in NES *versus* 5R/5A mutant **(Fig. 4d)**, the nuclear localization of ORF2, which is abrogated for the 5R/5A mutant, may affect pDC response against HEV. Since the 5R/5A mutant also does not produce infectious viruses **(Hervouet et al., 2022)**, pDC response against HEV might be modulated by viral particles release and/or nuclear ORF2.

IFNα production was lower when pDCs were co-cultured with PLC3 cells expressing the STOP mutant, as compared to cells expressing WT ORF2, suggesting that ORF2g/c forms contribute to the pDC IFN-I response in these cells. The ORF2g/c forms produced by the WT ORF2 may be recognized by the CLRs of pDCs, contributing to an enhanced response to immuno-stimulatory RNA. Alternatively, ORF2g/c expression may facilitate viral RNA transfer to pDC and its subsequent recognition by TLR7 in endosomes. However, the decrease of pDC response to the STOP mutant was not observed for HepG2/C3A cells **(Fig. 4e)**.

Here, we showed that the pDCs exhibited reduced IFNl7l response to HepG2/C3A expressing the 5R/5A mutant, and a complete absence of this response in PLC3 cells. This is likely due to differences in immune signaling among the two cell lines with the 5R/5A mutant. We observed that HEV infection impacted, in a cell-type dependent manner, the expression of *TNF* **(Figure 2a)**, a representative cytokine of NFKB signaling, known to modulate the expression of regulators of pDC adhesion and/or recruitment, including ICAMI **(Kim et al., 2008)** and various inflammatory chemokines **(Sedger & McDermott, 2014)**. We thus decided to analyze the effect of 5R/5A mutation on *TNF* expression in these two cell lines **(Supplementary figure 3a-b)**. We found that 5R/5A mutant significantly reduced *TNF* induction in PLC3 cells but not in HepG2/C3A cells as compared to HEV-WT, thus potentially explaining the cell type-specific regulation by ORF2i. Taken together, our results show that ORF2 protein expression and localization in producer cells modulate pDC response, with a magnitude that is cell-type dependent.

### ORF2 protein forms contribute to the robustness of contact between replicating cells and pDCs

We sought to determine whether HEV ORF2-mediated regulation of the strength and duration of cell-to-cell contacts could impact pDC response to infected cells. To achieve this, we developed a confocal imaging pipeline that quantifies cell proximity between pDCs and infected cells **(Supplementary Fig. 4a)**. Since more contrasting effects among ORF2 mutants were observed in PLC3 cells than in HepG2/C3A cells **(Fig. 4)**, we selected PLC3 cells for investigating the role of cell-to-cell contacts in pDC-mediated response. The pDCs and HEV-replicating cells were stained using CellTrace Violet (CTV) and CellTracker Red (CMPTX), respectively, and then co-cultured and fixed after 4 hours **(Fig. 5a)**. HEV-replicating cells were identified using the P3H2 anti-ORF2 antibody, which recognizes all forms of ORF2 (ORF2g/c/i) **(Bentaleb et al., 2022)**. The raw images were analyzed by an ImageJ macro-driven automatic analysis of cell-cell contacts. The two criteria for selection of contacts between pDCs and infected cells were the distance between the two cell types (1µm), and a contact area (>0µm^2^) between the surfaces of the two cell types **(Supplementary Fig. 4b)**.

**Figure 5.**
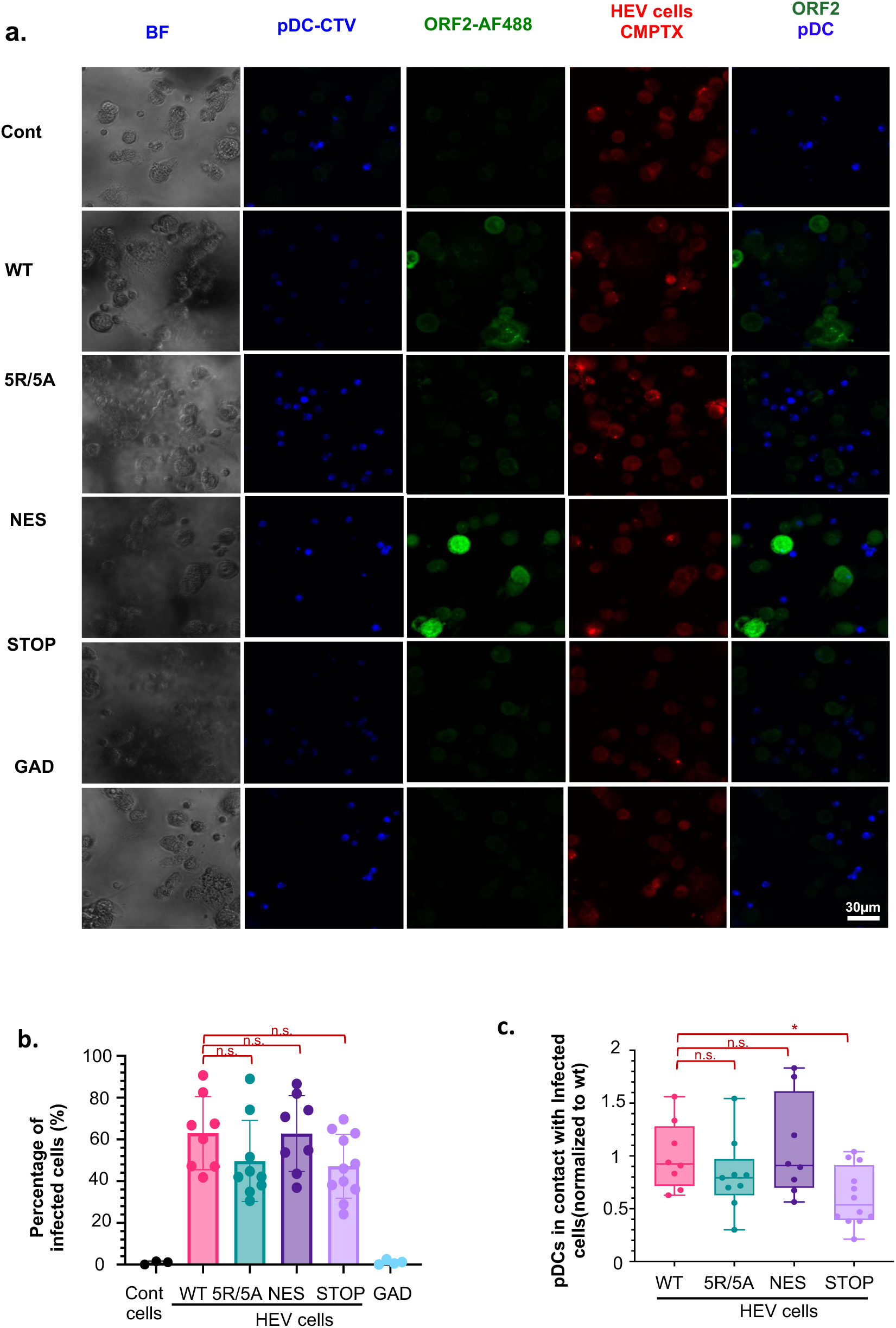
Effect of ORF2 expression and localization on physical contacts between HEV cells and pDCs, and pDC-mediated IFN response. (**A-E**) pDC co-cultured with HEV-replicating PLC3 cells 6 d.p.e, bearing the indicated mutations *versus* WT and GAD, as reference and negative control, respectively. Confocal imaging of pDCs and HEV co-culture performed after 4 hour-incubation. **(A)** Representative confocal stack-imaging of ORF2 immunostaining (green) in infected PLC3 cells, which were stained with Cell-Tracker Red prior co-cultures (CMTPX, red), combined with pDCs stained with Cell-Tracer Violet prior co-culture (CTV; blue). (**B**) Frequency of HEV-replicating (ORF2^+^) PLC3 cells detected among the cells defined as non-pDC (CTV^-^/CMTPX^+^); bars present means ± SD and each dot for independent image (n=4). (**C**) Contact/proximity of PLC3 infected cells and pDCs with detection of CTV/pDCs and CMTPX^+^ ORF2^+^/infected cells, as reference; bars present means ± SD and each dot for independent image (n[=[4). Statistical analysis was done using Wilcoxon rank sum test; *p* values as: * ≤0.05, ** ≤0.005 and *** ≤0.0005.

The percentage of ORF2 positive cells was within a comparable range among WT and the mutants in PLC3 cells **(Fig. 5b)**, in agreement with their similar RNA levels **(Fig. 4d)**. As compared to co-culture of pDCs with cells expressing WT ORF2, fewer cell-to-cell contacts **(Fig. 5c)** were observed between pDCs and cells expressing the 5R/5A (*i.e.*, reproducible but not significant) and a significant 40% decline in pDC contacts with cells expressing the STOP mutant. Taking into consideration that the STOP mutant displayed a reduced induction of pDC response (**Fig. 4f)**, this decline in pDC response to the STOP mutant could be due to reduced cell-to-cell contacts. Nevertheless, no difference was observed in the quality or strength of contacts, expressed in terms of contact area, across the mutants and WT **(Supplementary Fig. 4c)**. This indicates that pDC function is likely influenced by frequency of contact formation rather than the area of contact between the infected cell and pDC, which was found to be consistent across the wild-type and mutants.

Next, to further investigate the effect of ORF2 nuclear localization on immune signaling, we carried out single-cell imaging flow cytometry experiments in HepG2/C3A cells **(Figure 6a),** which had more robustly upregulated ISGs upon HEV infection independently of pDCs **(Figure 2a; left panel)**. Single-cell imaging flow cytometry allowed us to distinguish infected cells with (trORF2+) or without ORF2 nuclear translocation (trORF2-). We then examined these cells for IRF3 nuclear translocation, which is known to upregulate the expression of several cytokines, including IFN-I, IFN-III and CXCL10 **(Brownell et al., 2014)**, as well as surface molecules. Secretion of chemokines can lead to the recruitment of certain immune cell subsets. For instance, CXCL10 secretion triggers pDC recruitment **(Di Domizio et al., 2020)**.

**Figure 6.**
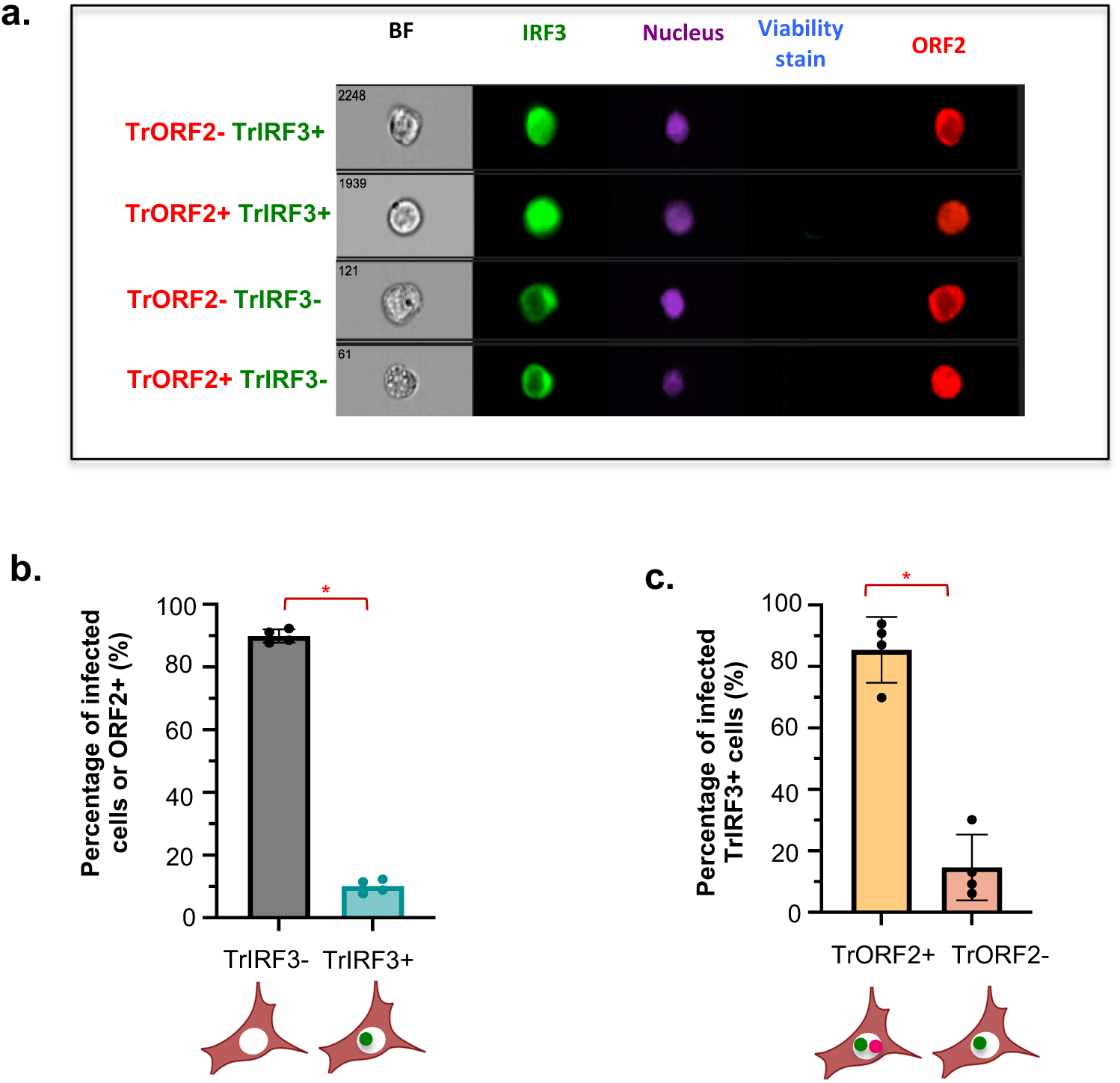
Single-cell analysis of IRF3 nuclear localization in HEV infected HepG2/C3A cells. HepG2/C3A cells expressing IRF3-GFP electroporated with WT HEV RNA and fixed 2 d.p.e. after staining with viability stain (Zombie aqua), and were then stained for ORF2 (APC) and nucleus (Hoechst). Single-cell images of viable ORF2+ cells were obtained and categorized according to the localization of IRF3 and ORF2. **(A)** Cells with translocation of ORF2 and IRF3 into the nucleus are represented by TrORF2+ and TrIRF3+. Cells without translocation of ORF2 and IRF3 into the nucleus are represented by TrORF2- and TrIRF3-**(B)** Percentage of HEV-replicating or ORF2+ cells with nuclear IRF3 translocation (ORF2+ TrIRF3+). **(C)** Percentage of ORF2 nuclear translocation (TrORF2+) among TrIRF3+ cells. Bars represent means ± SD and each dot represents one independent experiment (n=4). Statistical analysis using two-sided Wilcoxon rank sum test with continuity correction; *p* values as: * ≤0.05, ** ≤0.005 and *** ≤0.0005.

We observed that around 10% of trORF2+ cells displayed IRF3 nuclear translocation (TrIRF3+) **(Figure 6b)**. Around 80% of TrIRF3+ cells exhibited an ORF2 nuclear signal **(Figure 6c)**. These results revealed that the immune status of IRF3 in HepG2/C3A infected cells can differ depending on the nuclear translocation of HEV ORF2, further validating the impact of ORF2 nuclear localization on the immune status of the cells.

Together, our results suggest that ORF2g/c production likely modulates the number of cell-cell contacts and, subsequently, pDC-mediated IFN-I response. In the absence of ORF2g/c (STOP mutant), there is a significant decline in pDC-infected cell contacts. On the other hand, nuclear ORF2 can dictate the immune status in the infected cell *via* IRF3 signaling axis, possibly fueling the pDC response.

## DISCUSSION

Several studies have highlighted the suppressive impact of HEV on IFN response **(Devhare et al., 2021)**. Here, we investigated whether these inhibitory mechanisms can be overcome by pDCs, which act as immune sentinels against viral infections. Given the limited number of studies on cellular systems that effectively replicate HEV, we first determined the ability of three hepatoma cell lines to respond to several immune stimuli and replicate the viral genome. Based on their robust response to stimuli and their ability to replicate gt3 p6 strain HEV RNA, we selected PLC3 and HepG2/C3A cells for further analyses. Upon 6 days of HEV RNA electroporation, we observed a modest upregulation of IRF3 and NF-KB-mediated gene expression in HepG2/C3A cells. The genes that were significantly upregulated in HEV-replicating HepG2/C3A cells and included *ISG15*, *IFN-λ1*, *OAS-2* and *IL-6*. The upregulation of *IFN-λ1* expression in HepG2/C3A cells was in agreement with studies recognizing type III IFN response as a fundamental anti-HEV cytokine in hepatocytes both clinically **(Murata et al., 2020)** and *in vitro* **(Wu et al., 2018; Yin et al., 2017)**. Even though HEV can persist in the presence of a sustained type III IFN response **(Yin et al., 2017)**, this response prevents inflammation and helps maintain barrier integrity **(Broggi et al., 2020)**. Importantly, *TNF* upregulation was reduced by HEV in PLC3 cells, in agreement with previous data obtained in PLC3 cells **(Hervouet et al., 2022)**. This was not observed for HepG2/C3A cells, suggesting that HEV modulates NF-KB-mediated signaling in a cell-type specific manner. While exhibiting strong anti-viral effects **(Wang et al., 2016)**, TNFα is also implicated in pro-inflammatory processes exacerbating the severity of liver disease upon HEV infection **(Behrendt et al., 2017)**. Importantly, the differences in immune profiles of PLC3 cells and HepG2/C3A may be due to the expression of Hepatitis B surface antigen (HBsAg) by PLC3 cells, a known limitation of this cell type **(MacNab et al., 1976)**. Nevertheless, these cells may relevantly mimic the immune characteristics of individuals with pre-existing or previous infections or medical condition, predisposing them to HEV infection. In accordance - in the absence of pDC - secreted type III IFNs were undetectable in HEV-replicating PLC3 cells alone, and HEV-replicating cells also did not secrete any detectable IFNα, consistent with previous studies using HepG2 cells **(Yin et al., 2017)** and iPSC-HLCs **(Wu et al., 2018)**.

Upon co-culture of HEV-replicating HepG2/C3A cells with pDCs, ISGs as represented by *MXA*, *ISG15*, *OAS2* and *IFN-λ1*, were robustly upregulated. In fact, IFN-III response was also evident at the protein level after 48h of pDC co-culture with HEV-replicating HepG2/C3A cells. However, the expression of ISGs barely increased at 18h post co-culture with HEV-replicating PLC3 cells. Upon pDC co-culture of HEV-replicating PLC3 cells, only *IFN-λ1* was up-regulated significantly at mRNA level and, yet poorly detected as secreted protein, likely owing to the sensitivity limit of the currently available ELISA detection and thus requiring analysis with more sensitive techniques. IFN-λ1 expression by pDCs has been documented to actively participate in pDC response *e.g.*, against human cytomegalovirus and SARS-CoV-2 **(Venet et al., 2023; Yun et al., 2021)**. Whether the source of *IFN-λ1* upregulation were HEV-replicating cells, pDCs, or both cell types in concert, remains to be determined. Since the expression of these effectors is crucial for restricting viral replication **(Schneider et al., 2014)**, the absence of ISG upregulation presents a possible Achilles’ heel in host immune defense against HEV as this might contribute to HEV persistence in culture and its elimination might be challenging even after prolonged exposure to high doses of IFNs **(Todt et al., 2016)**. The lack of ISG expression was overcome by pDCs against HEV in the form of a strong IFN-I response when co-cultured with infected cells. This could activate ISG expression in neighboring cells **(Bourdon et al., 2020)** and also facilitate adaptive immune responses **(Crouse et al., 2015)**. Along the same line, the IFN-I response against HEV was dependent on pDC-infected cell contact formation. This was orchestrated by ICAMI and α_L_β_2_-integrin and TLR7, in agreement with results obtained for other viral infections **(Yin et al., 2017)**. The contact between pDCs and infected cells is beneficial for host defense, considering that HEV is resistant to exogenous IFN treatment **(Todt et al., 2016; Wang et al., 2016)**. On a different note, a prior study has shown that HEV-ORF3 can attenuate TLR7 expression **(Lei et al., 2018)**, this effect is less likely to be observed for pDCs as these are non-permissive to most viruses **(Reizis, 2019; Silvin et al., 2017; Venet et al., 2023)**. PDC response failed to control HEV replication after 18h of co-culture with PLC3 cells but effectively controlled HEV replication in HepG2/C3A under the same conditions. We observed more efficient or faster viral control in HepG2/C3A cells compared to PLC3 cells. This was likely due to a significant upregulation of ISGs (*ISG15, IFN-λ1, and OAS2*) - even in absence of pDCs - in HEV-replicating HepG2/C3A cells *versus* an inexistent one in HEV-replicating PLC3 cells. We hypothesize that the pre-existing ISG response in HEV-replicating HepG2/C3A, produced in the absence of pDCs, potentiates a more effective viral control in the presence of pDCs. Therefore, the initial ISG expression in infected cells may synergize the antiviral response by pDCs. Upon further scrutiny, pDC-mediated IFN response reduced HEV production by 50% 48h post co-culture in PLC3 cells. This also contributed to an overall decline in viral spread to neighboring cells. These results confirmed the need for a prolonged IFN response for controlling HEV replication.

In the absence of ORF2 nuclear translocation (5R/5A mutant), pDC IFN-I response was diminished in HepG2/C3A cells and abolished in PLC3 cells. Furthermore, when ORF2 accumulated in the nucleus (NES mutant), there was no difference in pDC response as compared to the wild-type. Additionally, the depletion of ORF2g/c secretion (STOP mutant) lowered the pDC response in PLC3 cells but not in HepG2/C3A cells. Altogether, these observations suggest that ORF2 nuclear translocation and secretion of glycosylated ORF2g/c forms might modulate pDC response in a cell-type dependent manner. Since many adhesion molecules are ISGs **(Ma et al., 2022; Parr & Parr, 2000)** and can impact the ability of HEV-mutant harboring cells to form cell-to-cell contact, we compared their pDC contact-forming propensity. Only cells harboring the STOP mutant differed from wild-type HEV cells in their ability to form contacts with pDCs. These results support the hypothesis of a reduced attraction of pDCs towards STOP-expressing PLC3 cells, as compared to control cells, due to the absence of ORF2g/c secretion. We did not see the same effect for HepG2/C3A cells, probably due to robust immune induction upon HEV infection. Alternative effector(s) favoring the contacts between pDCs and infected cells could be better expressed in HepG2/C3A cells compared to PLC3 cells, possibly due to the potent immune status, which may mask the effect of ORF2g/c on contact formation. However, this aspect requires further investigation. We should also take into account the decrease in viral infectivity in the absence of ORF2g/c **(Supplementary Fig. 2e)**, which could in turn lead to reduced pDC IFN response. Even though incubation with purified ORF2g/c does not impact HEV entry **(Yin et al., 2018)**, our results do raise further questions on whether its secretion can indirectly offer an advantage to HEV for spreading to the neighboring cell by improving cell-cell adhesion, a relatively easier route of viral transmission **(Zhong et al., 2013)**. Many studies have demonstrated that glycosylation is a common factor affecting cell-cell adhesion **(Gu et al., 2012; Ohtsubo & Marth, 2006)**. When rhesus macaques were infected with HEV variant that did not express glycosylated ORF2 forms, viral replication was attenuated and viral shedding in feces declined significantly as compared to infection with WT HEV **(Ralfs et al., 2023)**. Therefore, ORF2 g/c may not be essential for viral replication but it may be instrumental for efficient viral production.

Our findings emphasize the significance of the ongoing co-evolution between hosts and pathogens, allowing the virus to disseminate while concurrently enhancing its detection by immune cells. The difference in pDC response among HEV WT, NES *versus* 5R/5A still remains enigmatic but it paves a direction for further exploration in this exciting field. The presence of pDC response to 5R/5A expressing HepG2/C3A cells may imply regulation of pDC response at an additional level, *i.e.*, cell-type dependent regulation. Since pDCs mounted an IFN-I response against 5R/5A in HepG2/C3A cells, we propose that this detection could have been fueled by a successful upregulation of ISGs within HepG2/C3A cells compared to PLC3 cells upon HEV infection. Importantly, TNF signaling is induced in HepG2/C3A cells and inhibited in PLC3 cells. Since TNF signaling regulates cell adhesion via ICAM-1 **(Reglero-Real et al., 2014)** and activates an autocrine loop with low and sustained production of type I IFNs **(Yarilina et al., 2008)**, it could contribute to differential pDC response for the two cell-types. The difference in pDC-mediated IFN-I response to WT and 5R/5A mutant is particularly interesting and was observed for both cell lines, even though at different magnitudes. The 5R/5A mutant, which is characterized by an absence of nuclear ORF2, displayed a reduced or abolished pDC response *i.e.,* depending on the cell line. We then uncovered a cell line-specific induction of *TNF*, depending on ORF2 nuclear translocation. It indicates a possible differential modulation of downstream effector(s), including those involved in pDC cell adhesion/recruitment, and eventually leading to a weaker interferogenic synapse between PLC3 cells and pDCs, but not at play in HepG2/C3A cells. Further investigation will be needed to identify the effector(s) at play. We also investigated whether ORF2 nuclear localization could be involved in the immune state of the infected cells. We showed that IRF3 nuclear translocation tends to occur more in HEV-replicating cells with nuclear ORF2 than in HEV-replicating cells without nuclear ORF2. There could be several possible reasons for this activation of immune signaling, *i.e.*, intrusion of nuclear membrane, sensing of the viral ORF2 in the nucleus, or sensing of accidentally shuttled HEV RNA in the nucleus. It would be interesting to investigate this further. The activation of IRF3 signaling could, in turn, aid pDCs in viral sensing *via* cytokines and upregulation of other effector molecules.

Globally, the HEV mutants advanced our comprehension about viral regulation of the IFN response and also mechanistic nuances of viral recognition by pDCs. In summary, our findings suggest that pDC response is likely regulated by (i) transfer of viral entities to pDCs for recognition (ii) number of cell-to-cell contacts, and (iii) intrinsic immune state of the infected cells.

## Material and Methods

### Reagents and antibodies

Reagents used for pDC isolation are: Ficoll-Hypaque (GE Healthcare Life Sciences), BDCA-4-magnetic beads (MACS Miltenyi Biotec), LS Columns (MACS Miltenyi Biotec) and bottle top vacuum filters 0.22 μm (Nalgene). Other reagents included Poly(I:C) (LMW, Invivogen) as TLR3 agonist, Imiquimod as TLR7 agonist (Invivogen), and TLR7 antagonist (IRS661, 5’-TGCTTGCAAGCTTGCAAGCA-3’ synthesized on a phosphorothionate backbone MWG Biotech). The following antibodies were used: mouse anti-α_L_ integrin (clone 38; Antibodies Online); mouse anti-ICAM-1 (Clone LB-2; BD Bioscience); mouse anti-HEV ORF2 1E6, (Millipore, antibody registry #AB-827236), mouse anti-ORF2i/g/c P3H2 **(Bentaleb et al., 2022)** and mouse anti-ORF2i P1H1 **(Bentaleb et al., 2022)**. Goat anti-Mouse IgG2b Cross-Adsorbed Secondary Antibody Alexa Fluor 488 (Thermo Fisher Scientific Catalog #A-21141) were used for flow cytometry analysis.

### Cell Lines

PLC3 cells, a subclone of PLC3/PRF/5 cells **(Montpellier et al., 2018)**, Huh-7.5 cells (RRID:CVCL_7927) are hepatoma cells derived from Huh-7 cells **(Blight et al., 2002)**, and HepG2/C3A cells (ATCC CRL-3581) (provided by Dr. V.L. Dao Thi and Dr. D. Moradpour) were used. PLC3 and Huh-7.5 cells were maintained in Dulbecco’s modified Eagle medium (DMEM) supplemented with 10% FBS, 100 units (U)/ml penicillin, 100 mg/ml streptomycin, 2 mM L-glutamine and non-essential amino acids, 1 mM sodium pyruvate (Life Technologies) at 37**°**C/5% CO_2_. HepG2/C3A were maintained in DMEM (Glutamax, Pyr-) supplemented with 10% FBS, 100 units (U)/ml penicillin at 37**°**C/5% CO_2_.

### pDC isolation and culture

pDCs were isolated from blood from healthy adult human volunteers which was obtained from the ‘*Etablissement Francais du Sang’* (EFS; Auvergne Rhone Alpes, France) under the convention EFS 16–2066 and according to procedures approved by the EFS committee. Informed consent was obtained from all subjects in accordance with the Declaration of Helsinki. Information on sex and age was available for all subjects, yet we previously showed that pDC responses are within the same range for all donors **(Decembre et al., 2014)**. PBMCs were isolated using Ficoll-Hypaque density centrifugation and pDCs were positively selected from PBMCs using BDCA-4-magnetic beads (MACS Miltenyi Biotec), as previously described **(Decembre et al., 2014; Dreux et al., 2012; Venet et al., 2023)**. The typical yields of PBMCs and pDCs were 500-800×10^6^ and 1-2×10^6^ cells, respectively, with a typical purity of >95% pDCs. Isolated pDCs were maintained in RPMI 1640 medium (Life Technologies) supplemented with 10% FBS, 10 mM HEPES, 100 units/ml penicillin, 100 mg/ml streptomycin, 2 mM L-glutamine, non-essential amino acids and 1 mM sodium pyruvate at 37**°**C/5% CO_2_.

### Stimulation of cells for immune response using agonists and recombinant cytokines

PLC3 cells were seeded in 24-well plates at a concentration of 8.10^4^ cells/well. After 24h, cells were incubated with poly(I:C) LMW in complete medium at a concentration of 25 μg/ml and 100 μg/ml, in a total volume of 0.5ml medium. Alternatively, cells were transfected with poly(I:C) LMW at a concentration of 0.3 μg/ml and 2 μg/ml. Lipofectamine 2000 (Invitrogen) in Opti-MEM I reduced serum media was used for transfection according to the manufacturer’s protocol. IFN-β subtype 1a (Invitrogen, Catalog#PHC4244), IFN-λ3/Interleukin-28B (IL-28B) (PBL Interferon Source #11820-1), TNF (Cell Signalling Technology #16769) were added at a concentration of 100U/ml, 100ng/mL and 535 U/mL, respectively, in a total volume of 0.5 mL medium per well. The cells were harvested 6h post-treatment for RNA extraction.

### Analysis of transcriptional levels by RT-qPCR

RNAs were isolated from cells harvested in guanidinium thiocyanate citrate buffer (GTC) by phenol/chloroform extraction procedure, as described previously (Assil, Coléon, et al., 2019). The mRNA levels of human *MXA, ISG15*, *IFN-λ1, IL6, TNF, ORF2* and glyceraldehyde-3-phosphate dehydrogenase (*GADPH*) were determined by RT-qPCR using High-Capacity cDNA Reverse Transcription Kit (ref# 4368813) and PowerUp™ SYBR™ Green Master Mix (ref# A25778) for qPCR run on QuantStudio6 PCR system and analyzed using QuantStudio Design and Analysis Software (Thermo Fisher Scientific). The mRNA levels were normalized to *GADPH* mRNA levels. The sequences of the primers used for RT-qPCR are as listed in **supplementary Table 1**.

### Plasmids and electroporation for HEV infectious system

The plasmid pBlueScript SK(+) carrying the DNA of the full length genome of adapted gt3 Kernow C-1 p6 strain, (GenBank accession number JQ679013, kindly provided by Dr S.U. Emerson), was used as the wild-type genome. The 5R/5A and NES mutant plasmids were previously described **(Hervouet et al., 2022)**. The STOP mutant was generated by site directed mutagenesis using the following primers :

P6/mutstop-F : TGTTCTGCTGCTGTagTTCGTGTTTCTGCCTATGC

P6/mutstop-R : GGCAGAAACACGAActACAGCAGCAGAACAACCC

The Stop mutation was introduced by sequential PCR steps, as previously described **(Ankavay et al., 2019)**, and verified by DNA sequencing. The non-replicative GAD mutant plasmid was a gift from Dr V. L. Dao Thi **(Emerson et al., 2013)**. To prepare genomic HEV RNAs (capped RNA), the wild type and mutant pBlueScript SK(+) HEV plasmids were linearized at their 3′ end with the MluI restriction enzyme (NEB) and transcribed with the mMESSAGE mMACHINE T7 Transcription kit (Ambion #1344). 10 μg capped RNAs were next delivered to cells by electroporation using a Gene Pulser X cell 617BR 11218 (BioRad).

### Co-culture of infected cells with isolated pDCs and ELISA

Unless otherwise indicated, 5×10^4^ pDCs were co-cultured with 6×10^4^ PLC3 or HepG2/C3A cells, electroporated or not, or as a control condition, filtered supernatants (0.4 µm). Infected cells were electroporated for 48, 72 or 144 hours (as indicated) in a 200 μL final volume in 96-well round-bottom plates incubated at 37**°**C/5% CO_2_. When indicated, cells were co-cultured in 96-well format transwell chambers (Corning) with a 0.4 µm permeable membrane. Sixteen to eighteen hours later (as specified), cell-culture supernatants were collected and the levels of IFNα were measured using a commercially available specific ELISA kit (PBL Interferon Source Catalog# 411052) with an assay range of 12.5-500 pg/ml.

### Flow cytometry-based viral spread assays and imaging flow cytometry by ImageStream X Technology

PLC3 cells were transduced with lentiviral-based vector pseudotyped with vesicular stomatitis virus (VSV) glycoprotein to stably express GFP. After immuno-isolation, pDCs were stained for 20 minutes at 37°C in the dark. Labeled pDCs were then spun down and re-suspended in pDC culture medium. 5×10^4^ pDCs were co-cultured with 3×10^4^ HEV-replicating cells (electroporated 6 days prior to co-culture) and with 3×10^4^ GFP^+^ uninfected cells for 48 hours at 37°C/5% CO_2_. The level of viral spread from HEV-replicating cells (ORF2^+^) to uninfected cells (GFP^+^) during co-culture was determined by flow cytometric analysis as the frequency of infected cells (ORF2^+^/GFP^+^ population) among the GFP^+^ cell population and similarly in GFP^-^ populations. At the indicated times, harvested cells were resuspended using 0.48 mM EDTA-PBS solution for the co-culture with pDCs. Cells were then incubated with 1 μL/mL viability marker diluted in PBS for 20 minutes at RT. Cytoperm/Cytofix^TM^ and permeabilization-wash solutions (BD Bioscience) were used in the subsequent stages of the ORF2 staining protocol optimized in-house for flow cytometry. Cells were fixed with Cytofix for 20 minutes at 4°C and were then washed twice and resuspended in 1x Permwash. The fixed cells were treated with cold methanol (−20°C) for 45 minutes and washed twice with 1x Permwash. The cells were then incubated with 1E6 anti-HEV antibody for a minimum of 3 hours. After another wash, cells were incubated with a secondary antibody, *i.e.*, Goat anti-Mouse IgG2b Cross-Adsorbed Secondary Antibody Alexa Fluor 488 (Thermo Fisher Scientific Catalog #A-21141) for 2 hours. The samples were acquired after final two washes. Flow cytometer analysis was performed using a Canto II Becton Dickinson using BD FACSDIVA v8.1 software and the data were analyzed using Flow Jo 10.8.1 software (Tree Star). CellTrace Violet (CTV) Cell Proliferation kit (Life Technologies) was used to stain pDCs. LIVE/DEAD™ Fixable Near-IR - Dead Cell Stain Kit (Thermo Fisher Scientific ref #L10119) was used to check viability. For imaging flow cytometry, cells expressing IRF3-GFP (construct was kindly provided by Dr. Marco Binder) **(Willemsen et al., 2017)** were electroporated with HEV WT RNA. The cells were then harvested at 48h.p.e., followed by viability staining with Zombie Aqua (Biolegend, Cat # 423101), and then ORF2 staining protocol. At the final step, Hoechst (Thermo Fisher Scientific ref #H1399) was used to stain the nuclei of the cells. After acquisition by ImageStreamX MarkII (Amnis - Millipore), analysis was done using IDEAS software.

### Microscopy and analysis of images

At 4 hours post-co-culture, CTV-stained-pDCs and infected PLC3 and HepG2/C3A cells were fixed with PFA 4%, followed by immunostaining using anti-ORF2 P3H2 antibodies with a protocol optimized previously **(Bentaleb et al., 2022)**. Confocal imaging was performed using Zeiss LSM980 scanning confocal microscopy. Co-cultured cells were automatically segmented based on the CMPTx staining for cell line used and CTV labeling for pDCs. Next, the infected cells were identified by ORF2 expression, and number/frequency of infected cells were quantified using one of two similar versions of a home-made ImageJ macro (https://github.com/jbrocardplatim/PDC-contacts), adapted to HepG2/C3A or PLC3 cells. The frequency of pDCs within 3 microns of any cell was calculated, as well as the number of pDCs within 3 microns of any infected cell (from 0 to 3+). An additional macro was also used to measure the frequency of pDCs closer than 1 micron to infected cells and/or in direct contact; the individual contact areas (in µm²) were measured as well (https://github.com/jbrocardplatim/PDC-contacts). The calculation formula for contact frequencies considered variations in the number of infected cells to mitigate potential biases.

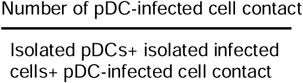

To compare the ORF2 localization in mutants and wild-type bearing cells, nuclei were stained with NucSpot® Live 650 (Biotium, Catalog#40082) and HEV-ORF2 was stained with 1E6 antibody. For all microscopy experiments, cells were seeded in 96-Well Optical-Bottom Plate (Thermo Fisher Scientific), coated with poly-L-lysine (P6282, Sigma-Aldrich).

### Western blotting and immunoprecipitation analyses

Western blotting (WB) and immunoprecipitation (IP) analyses were performed as described previously **(Bentaleb et al., 2022).** For WB, supernatants and lysates of WT and STOP expressing PLC3 cells were heated for 20 min at 80 °C in the presence of reducing Laemmli buffer. Samples were then separated by 10% SDS-PAGE and transferred onto nitrocellulose membranes (Hybond-ECL, Amersham). ORF2 proteins were detected with 1E6 antibody and peroxidase-conjugated anti-mouse antibodies. The detection of proteins was done by chemiluminescence analysis (ECL, Amersham). For IP, P3H2 and P1H1 antibodies were bound to M-280 Dynabeads (Thermofisher) overnight at 37°C following the manufacturer’s recommendations. Beads were washed and then incubated for 1h at room temperature with supernatants (heat-inactivated) or cell lysates. Beads were washed and then heated at 80°C for 20 min in reducing Laemmli buffer. ORF2 proteins were detected by WB using the 1E6 antibody.

### Statistical analysis

Statistical analysis was performed using R software environment for statistical computing and graphics (version 3.3.2). For quantifications by ELISA, RT-qPCR, and flow cytometry, analyses were performed using Wilcoxon rank-sum exact test and P value adjustment method: Bonferroni or FDR was applied when mentioned. The figures were prepared using PRISM software (version 10.2.1).

## Acknowledgements

We would like to thank ANRS-MIE for providing financial support (ANRS-MIE grant number: AAP2020-2/ECTZ133955) and ‘Contrats doctoraux Lyon 1 dédiés à l’International’ from Université Lyon 1) for GJ’s PhD fellowship. This work was also supported by grants from the Agence Nationale de la Recherche (ANRJCJC-iSYN); the Agence Nationale pour la Recherche contre le SIDA et les Hépatites Virales (ANRS – N21006CR and N19017CR); the UDL/ANR IA ELAN ERC (G19005CC) to MD. We acknowledge Dr. Julie Lucifora, Dr. David Durantel, Dr. Elena Tomasello, Dr. Alexandre Belot, Dr. Søren Riis Paludan, Dr. Marco Binder and Dr. Antoine Marçais for scientific discussions. We thank SFR Biosciences (UMS3444/CNRS, US8/Inserm, ENS de Lyon, UCBL) including the PLATIM and AniRA-cytometry facilities, for technical assistance in imaging and FACS analyses, respectively. Special thanks to Dr. Marion Delphin, Dr. Vladimir Goncalves Magalhaes and Roxanne Fouille for their contribution in supporting experiments. The contribution of the EFS Confluence/Decine-Lyon is also noteworthy. Finally, we thank current and former members of the VIV team for helpful discussions and support: Dr. Margarida Sa Ribeiro, Célia Nuovo, Matteo Agostini, and Manon Venet.

## References

1. Andonov, A., Robbins, M., Borlang, J., Cao, J., Hatchette, T., Stueck, A., Deschambault, Y., Murnaghan, K., Varga, J., & Johnston, L. (2019). Rat Hepatitis E Virus Linked to Severe Acute Hepatitis in an Immunocompetent Patient. The Journal of Infectious Diseases, 220(6), 951–955. 10.1093/infdis/jiz025

2. Ankavay, M., Montpellier, C., Sayed, I. M., Saliou, J. M., Wychowski, C., Saas, L., Duvet, S., Aliouat-Denis, C. M., Farhat, R., de Masson d’Autume, V., Meuleman, P., Dubuisson, J., & Cocquerel, L. (2019). New insights into the ORF2 capsid protein, a key player of the hepatitis E virus lifecycle. Scientific Reports. 10.1038/s41598-019-42737-2

3. Assil, S., Coléon, S., Dong, C., Décembre, E., Sherry, L., Allatif, O., Webster, B., & Dreux, M. (2019). Plasmacytoid Dendritic Cells and Infected Cells Form an Interferogenic Synapse Required for Antiviral Responses. Cell Host and Microbe. 10.1016/j.chom.2019.03.005

4. Assil, S., Futsch, N., Décembre, E., Alais, S., Gessain, A., Cosset, F.-L., Mahieux, R., Dreux, M., & Dutartre, H. (2019). Sensing of cell-associated HTLV by plasmacytoid dendritic cells is regulated by dense β-galactoside glycosylation. PLOS Pathogens, 15(2), e1007589. 10.1371/journal.ppat.1007589

5. Bagdassarian, E., Doceul, V., Pellerin, M., Demange, A., Meyer, L., Jouvenet, N., & Pavio, N. (2018). The Amino-Terminal Region of Hepatitis E Virus ORF1 Containing a Methyltransferase (Met) and a Papain-Like Cysteine Protease (PCP) Domain Counteracts Type I Interferon Response. Viruses, 10(12), 726. 10.3390/v10120726

6. Behrendt, P., Lüth, S., Dammermann, W., Drave, S., Brown, R. J. P., Todt, D., Schnoor, U., Steinmann, E., Wedemeyer, H., Pischke, S., & Iking-Konert, C. (2017). Exacerbation of hepatitis E virus infection during anti-TNFα treatment. Joint Bone Spine, 84(2), 217–219. 10.1016/j.jbspin.2016.09.017

7. Bentaleb, C., Hervouet, K., Montpellier, C., Camuzet, C., Ferrié, M., Burlaud-Gaillard, J., Bressanelli, S., Metzger, K., Werkmeister, E., Ankavay, M., Janampa, N. L., Marlet, J., Roux, J., Deffaud, C., Goffard, A., Rouillé, Y., Dubuisson, J., Roingeard, P., Aliouat-Denis, C.-M., & Cocquerel, L. (2022). The endocytic recycling compartment serves as a viral factory for hepatitis E virus. Cellular and Molecular Life Sciences, 79(12), 615. 10.1007/s00018-022-04646-y

8. Bermejo-Jambrina, M., Eder, J., Helgers, L. C., Hertoghs, N., Nijmeijer, B. M., Stunnenberg, M., & Geijtenbeek, T. B. H. (2018). C-Type Lectin Receptors in Antiviral Immunity and Viral Escape. Frontiers in Immunology, 9, 590. 10.3389/fimmu.2018.00590

9. Blight, K. J., McKeating, J. A., & Rice, C. M. (2002). Highly Permissive Cell Lines for Subgenomic and Genomic Hepatitis C Virus RNA Replication. Journal of Virology, 76(24), 13001–13014. 10.1128/JVI.76.24.13001-13014.2002

10. Bourdon, M., Manet, C., & Montagutelli, X. (2020). Host genetic susceptibility to viral infections: The role of type I interferon induction. Genes & Immunity, 21(6–8), 365–379. 10.1038/s41435-020-00116-2

11. Broggi, A., Granucci, F., & Zanoni, I. (2020). Type III interferons: Balancing tissue tolerance and resistance to pathogen invasion. Journal of Experimental Medicine, 217(1), e20190295. 10.1084/jem.20190295

12. Brownell, J., Bruckner, J., Wagoner, J., Thomas, E., Loo, Y.-M., Gale, M., Liang, T. J., & Polyak, S. J. (2014). Direct, Interferon-Independent Activation of the CXCL10 Promoter by NF-κB and Interferon Regulatory Factor 3 during Hepatitis C Virus Infection. Journal of Virology, 88(3), 1582–1590. 10.1128/JVI.02007-13

13. Cambi, A., Koopman, M., & Figdor, C. G. (2005). How C-type lectins detect pathogens: C-type lectins and pathogens. Cellular Microbiology, 7(4), 481–488. 10.1111/j.1462-5822.2005.00506.x

14. Cervantes-Barragan, L., Lewis, K. L., Firner, S., Thiel, V., Hugues, S., Reith, W., Ludewig, B., & Reizis, B. (2012). Plasmacytoid dendritic cells control T-cell response to chronic viral infection. Proceedings of the National Academy of Sciences of the United States of America. 10.1073/pnas.1117359109

15. Crouse, J., Kalinke, U., & Oxenius, A. (2015). Regulation of antiviral T cell responses by type I interferons. Nature Reviews Immunology, 15(4), 231–242. 10.1038/nri3806

16. Devhare, P., Madiyal, M., Mukhopadhyay, C., Shetty, S., & Shastry, S. (2021). Interplay between Hepatitis E Virus and Host Cell Pattern Recognition Receptors. International Journal of Molecular Sciences, 22(17), 9259. 10.3390/ijms22179259

17. Di Domizio, J., Belkhodja, C., Chenuet, P., Fries, A., Murray, T., Mondéjar, P. M., Demaria, O., Conrad, C., Homey, B., Werner, S., Speiser, D. E., Ryffel, B., & Gilliet, M. (2020). The commensal skin microbiota triggers type I IFN–dependent innate repair responses in injured skin. Nature Immunology, 21(9), 1034–1045. 10.1038/s41590-020-0721-6

18. Doceul, V., Bagdassarian, E., Demange, A., & Pavio, N. (2016). Zoonotic Hepatitis E Virus: Classification, Animal Reservoirs and Transmission Routes. Viruses, 8(10), 270. 10.3390/v8100270

19. Dong, C., Zafrullah, M., Mixson-Hayden, T., Dai, X., Liang, J., Meng, J., & Kamili, S. (2012). Suppression of interferon-α signaling by hepatitis E virus. Hepatology, 55(5), 1324–1332. 10.1002/hep.25530

20. Doyle, E. H., Rahman, A., Aloman, C., Klepper, A. L., El-Shamy, A., Eng, F., Rocha, C., Kim, S., Haydel, B., Florman, S. S., Fiel, M. I., Schiano, T., & Branch, A. D. (2019). Individual liver plasmacytoid dendritic cells are capable of producing IFNα and multiple additional cytokines during chronic HCV infection. PLOS Pathogens, 15(7), e1007935. 10.1371/journal.ppat.1007935

21. Dreux, M., Garaigorta, U., Boyd, B., Décembre, E., Chung, J., Whitten-Bauer, C., Wieland, S., & Chisari, F. V. (2012). Short-Range Exosomal Transfer of Viral RNA from Infected Cells to Plasmacytoid Dendritic Cells Triggers Innate Immunity. Cell Host & Microbe, 12(4), 558–570. 10.1016/j.chom.2012.08.010

22. Emerson, S. U., Nguyen, H. T., Torian, U., Mather, K., & Firth, A. E. (2013). An essential RNA element resides in a central region of hepatitis E virus ORF2. Journal of General Virology, 94(7), 1468–1476. 10.1099/vir.0.051870-0

23. Florentin, J., Aouar, B., Dental, C., Thumann, C., Firaguay, G., Gondois-Rey, F., Soumelis, V., Baumert, T. F., Nunès, J. A., Olive, D., Hirsch, I., & Stranska, R. (2012). HCV glycoprotein E2 is a novel BDCA-2 ligand and acts as an inhibitor of IFN production by plasmacytoid dendritic cells. Blood, 120(23), 4544–4551. 10.1182/blood-2012-02-413286

24. Gibson, S. J., Lindh, J. M., Riter, T. R., Gleason, R. M., Rogers, L. M., Fuller, A. E., Oesterich, J. L., Gorden, K. B., Qiu, X., McKane, S. W., Noelle, R. J., Miller, R. L., Kedl, R. M., Fitzgerald-Bocarsly, P., Tomai, M. A., & Vasilakos, J. P. (2002). Plasmacytoid dendritic cells produce cytokines and mature in response to the TLR7 agonists, imiquimod and resiquimod. Cellular Immunology, 218(1–2), 74–86. 10.1016/S0008-8749(02)00517-8

25. Gilliet, M., Cao, W., & Liu, Y.-J. (2008). Plasmacytoid dendritic cells: Sensing nucleic acids in viral infection and autoimmune diseases. Nature Reviews Immunology, 8(8), 594–606. 10.1038/nri2358

26. Graff, J., Torian, U., Nguyen, H., & Emerson, S. U. (2006). A Bicistronic Subgenomic mRNA Encodes both the ORF2 and ORF3 Proteins of Hepatitis E Virus. Journal of Virology, 80(12), 5919–5926. 10.1128/JVI.00046-06

27. Grange, Z. L., Goldstein, T., Johnson, C. K., Anthony, S., Gilardi, K., Daszak, P., Olival, K. J., O’Rourke, T., Murray, S., Olson, S. H., Togami, E., Vidal, G., Expert Panel, PREDICT Consortium, Mazet, J. A. K., Anderson, K., Auewarakul, P., Coffey, L., Corley, R.,… Fine, A. (2021). Ranking the risk of animal-to-human spillover for newly discovered viruses. Proceedings of the National Academy of Sciences, 118(15), e2002324118. 10.1073/pnas.2002324118

28. Gu, J., Isaji, T., Xu, Q., Kariya, Y., Gu, W., Fukuda, T., & Du, Y. (2012). Potential roles of N-glycosylation in cell adhesion. Glycoconjugate Journal, 29(8–9), 599–607. 10.1007/s10719-012-9386-1

29. Hervouet, K., Ferrié, M., Ankavay, M., Montpellier, C., Camuzet, C., Alexandre, V., Dembélé, A., Lecoeur, C., Foe, A. T., Bouquet, P., Hot, D., Vausselin, T., Saliou, J.-M., Salomé-Desnoulez, S., Vandeputte, A., Marsollier, L., Brodin, P., Dreux, M., Rouillé, Y.,… Cocquerel, L. (2022). An Arginine-Rich Motif in the ORF2 capsid protein regulates the hepatitis E virus lifecycle and interactions with the host cell. PLOS Pathogens, 18(8), e1010798. 10.1371/journal.ppat.1010798

30. Kim, H., Hwang, J.-S., Woo, C.-H., Kim, E.-Y., Kim, T.-H., Cho, K.-J., Seo, J.-M., Lee, S.-S., & Kim, J.-H. (2008). TNF-α-induced up-regulation of intercellular adhesion molecule-1 is regulated by a Rac-ROS-dependent cascade in human airway epithelial cells. Experimental and Molecular Medicine, 40(2), 167. 10.3858/emm.2008.40.2.167

31. Knowles, B. B., Howe, C. C., & Aden, D. P. (1980). Human Hepatocellular Carcinoma Cell Lines Secrete the Major Plasma Proteins and Hepatitis B Surface Antigen. Science, 209(4455), 497–499. 10.1126/science.6248960

32. Koonin, E. V., Gorbalenya, A. E., Purdy, M. A., Rozanov, M. N., Reyes, G. R., & Bradley, D. W. (1992). Computer-assisted assignment of functional domains in the nonstructural polyprotein of hepatitis E virus: Delineation of an additional group of positive-strand RNA plant and animal viruses. Proceedings of the National Academy of Sciences, 89(17), 8259–8263. 10.1073/pnas.89.17.8259

33. Lazear, H. M., Schoggins, J. W., & Diamond, M. S. (2019). Shared and Distinct Functions of Type I and Type III Interferons. Immunity, 50(4), 907–923. 10.1016/j.immuni.2019.03.025

34. Lee, G.-H., Tan, B.-H., Chi-Yuan Teo, E., Lim, S.-G., Dan, Y.-Y., Wee, A., Kim Aw, P. P., Zhu, Y., Hibberd, M. L., Tan, C.-K., Purdy, M. A., & Teo, C.-G. (2016). Chronic Infection With Camelid Hepatitis E Virus in a Liver Transplant Recipient Who Regularly Consumes Camel Meat and Milk. Gastroenterology, 150(2), 355–357.e3. 10.1053/j.gastro.2015.10.048

35. Lei, Q., Li, L., Zhang, S., Li, T., Zhang, X., Ding, X., & Qin, B. (2018). HEV ORF3 downregulates TLR7 to inhibit the generation of type I interferon via impairment of multiple signaling pathways. Scientific Reports, 8(1), 8585. 10.1038/s41598-018-26975-4

36. Lenggenhager, D., Gouttenoire, J., Malehmir, M., Bawohl, M., Honcharova-Biletska, H., Kreutzer, S., Semela, D., Neuweiler, J., Hürlimann, S., Aepli, P., Fraga, M., Sahli, R., Terracciano, L., Rubbia-Brandt, L., Müllhaupt, B., Sempoux, C., Moradpour, D., & Weber, A. (2017). Visualization of hepatitis E virus RNA and proteins in the human liver. Journal of Hepatology, 67(3), 471–479. 10.1016/j.jhep.2017.04.002

37. Lhomme, S., Marion, O., Abravanel, F., Izopet, J., & Kamar, N. (2020). Clinical Manifestations, Pathogenesis and Treatment of Hepatitis E Virus Infections. Journal of Clinical Medicine, 9(2), 331. 10.3390/jcm9020331

38. Li, P., Liu, J., Li, Y., Su, J., Ma, Z., Bramer, W. M., Cao, W., De Man, R. A., Peppelenbosch, M. P., & Pan, Q. (2020). The global epidemiology of hepatitis E virus infection: A systematic review and meta□analysis. Liver International, 40(7), 1516– 1528. 10.1111/liv.14468

39. Lin, S., Yang, Y., Nan, Y., Ma, Z., Yang, L., & Zhang, Y.-J. (2019). The Capsid Protein of Hepatitis E Virus Inhibits Interferon Induction via Its N-Terminal Arginine-Rich Motif. Viruses, 11(11), 1050. 10.3390/v11111050

40. Ma, N., Lu, J., Pei, Y., & Robertson, E. S. (2022). Transcriptome reprogramming of Epstein-Barr virus infected epithelial and B cells reveals distinct host-virus interaction profiles. Cell Death & Disease, 13(10), 894. 10.1038/s41419-022-05327-1

41. Ma, Z., De Man, R. A., Kamar, N., & Pan, Q. (2022). Chronic hepatitis E: Advancing research and patient care. Journal of Hepatology, 77(4), 1109–1123. 10.1016/j.jhep.2022.05.006

42. MacNab, G. M., Alexander, J. J., Lecatsas, G., Bey, E. M., & Urbanowicz, J. M. (1976). Hepatitis B surface antigen produced by a human hepatoma cell line. British Journal of Cancer, 34(5), 509–515. 10.1038/bjc.1976.205

43. Marlin, S. D., & Springer, T. A. (1987). Purified intercellular adhesion molecule-1 (ICAM-1) is a ligand for lymphocyte function-associated antigen 1 (LFA-1). Cell, 51(5), 813–819. 10.1016/0092-8674(87)90104-8

44. Meyer-Wentrup, F., Benitez-Ribas, D., Tacken, P. J., Punt, C. J. A., Figdor, C. G., De Vries, I. J. M., & Adema, G. J. (2008). Targeting DCIR on human plasmacytoid dendritic cells results in antigen presentation and inhibits IFN-α production. Blood, 111(8), 4245–4253. 10.1182/blood-2007-03-081398

45. Moal, V., Textoris, J., Ben Amara, A., Mehraj, V., Berland, Y., Colson, P., & Mege, J.-L. (2013). Chronic Hepatitis E Virus Infection Is Specifically Associated With an Interferon-Related Transcriptional Program. The Journal of Infectious Diseases, 207(1), 125–132. 10.1093/infdis/jis632

46. Montpellier, C., Wychowski, C., Sayed, I. M., Meunier, J.-C., Saliou, J.-M., Ankavay, M., Bull, A., Pillez, A., Abravanel, F., Helle, F., Brochot, E., Drobecq, H., Farhat, R., Aliouat-Denis, C.-M., Haddad, J. G., Izopet, J., Meuleman, P., Goffard, A., Dubuisson, J., & Cocquerel, L. (2018). Hepatitis E Virus Lifecycle and Identification of 3 Forms of the ORF2 Capsid Protein. Gastroenterology, 154(1), 211–223.e8. 10.1053/j.gastro.2017.09.020

47. Murata, K., Kang, J.-H., Nagashima, S., Matsui, T., Karino, Y., Yamamoto, Y., Atarashi, T., Oohara, M., Uebayashi, M., Sakata, H., Matsubayashi, K., Takahashi, K., Arai, M., Mishiro, S., Sugiyama, M., Mizokami, M., & Okamoto, H. (2020). IFN-λ3 as a host immune response in acute hepatitis E virus infection. Cytokine, 125, 154816. 10.1016/j.cyto.2019.154816

48. Nagashima, S., Primadharsini, P. P., Nishiyama, T., Takahashi, M., Murata, K., & Okamoto, H. (2023). Development of a HiBiT-tagged reporter hepatitis E virus and its utility as an antiviral drug screening platform. Journal of Virology, 97(9), e00508–23. 10.1128/jvi.00508-23

49. Nan, Y., Yu, Y., Ma, Z., Khattar, S. K., Fredericksen, B., & Zhang, Y.-J. (2014). Hepatitis E Virus Inhibits Type I Interferon Induction by ORF1 Products. Journal of Virology. 10.1128/jvi.01935-14

50. Nimgaonkar, I., Ding, Q., Schwartz, R. E., & Ploss, A. (2018). Hepatitis E virus: Advances and challenges. Nature Reviews Gastroenterology & Hepatology, 15(2), 96–110. 10.1038/nrgastro.2017.150

51. Ohtsubo, K., & Marth, J. D. (2006). Glycosylation in Cellular Mechanisms of Health and Disease. Cell, 126(5), 855–867. 10.1016/j.cell.2006.08.019

52. Parr, M. B., & Parr, E. L. (2000). Interferon□γ up□regulates intercellular adhesion molecule□1 and vascular cell adhesion molecule□1 and recruits lymphocytes into the vagina of immune mice challenged with herpes simplex virus□2. Immunology, 99(4), 540–545. 10.1046/j.1365-2567.2000.00980.x

53. Ralfs, P., Holland, B., Salinas, E., Bremer, B., Wang, M., Zhu, J., Ambardekar, C., Blasczyk, H., Walker, C. M., Feng, Z., & Grakoui, A. (2023). Soluble ORF2 protein enhances HEV replication and induces long-lasting antibody response and protective immunity in vivo. Hepatology, 78(6), 1867–1881. 10.1097/HEP.0000000000000421

54. Reglero-Real, N., Álvarez-Varela, A., Cernuda-Morollón, E., Feito, J., Marcos-Ramiro, B., Fernández-Martín, L., Gómez-Lechón, M. J., Muntané, J., Sandoval, P., Majano, P. L., Correas, I., Alonso, M. A., & Millán, J. (2014). Apicobasal Polarity Controls Lymphocyte Adhesion to Hepatic Epithelial Cells. Cell Reports, 8(6), 1879–1893. 10.1016/j.celrep.2014.08.007

55. Rehwinkel, J., & Gack, M. U. (2020). RIG-I-like receptors: Their regulation and roles in RNA sensing. Nature Reviews Immunology, 20(9), 537–551. 10.1038/s41577-020-0288-3

56. Reizis, B. (2019). Plasmacytoid Dendritic Cells: Development, Regulation, and Function. Immunity, 50(1), 37–50. 10.1016/j.immuni.2018.12.027

57. Sari, G., Mulders, C. E., Zhu, J., Van Oord, G. W., Feng, Z., Kreeft□Voermans, J. J. C., Boonstra, A., & Vanwolleghem, T. (2021). Treatment induced clearance of hepatitis E viruses by interferon□lambda in liver□humanized mice. Liver International, 41(12), 2866–2873. 10.1111/liv.15033

58. Sayed, I. M., Verhoye, L., Cocquerel, L., Abravanel, F., Foquet, L., Montpellier, C., Debing, Y., Farhoudi, A., Wychowski, C., Dubuisson, J., Leroux-Roels, G., Neyts, J., Izopet, J., Michiels, T., & Meuleman, P. (2017). Study of hepatitis E virus infection of genotype 1 and 3 in mice with humanised liver. Gut, 66(5), 920–929. 10.1136/gutjnl-2015-311109

59. Schemmerer, M., Apelt, S., Trojnar, E., Ulrich, R. G., Wenzel, J. J., & Johne, R. (2016). Enhanced replication of hepatitis E virus strain 47832C in an A549-derived subclonal cell line. Viruses. 10.3390/v8100267

60. Schneider, W. M., Chevillotte, M. D., & Rice, C. M. (2014). Interferon-Stimulated Genes: A Complex Web of Host Defenses. Annual Review of Immunology, 32(1), 513–545. 10.1146/annurev-immunol-032713-120231

61. Sedger, L. M., & McDermott, M. F. (2014). TNF and TNF-receptors: From mediators of cell death and inflammation to therapeutic giants – past, present and future. Cytokine & Growth Factor Reviews, 25(4), 453–472. 10.1016/j.cytogfr.2014.07.016

62. Shukla, P., Nguyen, H. T., Faulk, K., Mather, K., Torian, U., Engle, R. E., & Emerson, S. U. (2012). Adaptation of a Genotype 3 Hepatitis E Virus to Efficient Growth in Cell Culture Depends on an Inserted Human Gene Segment Acquired by Recombination. Journal of Virology, 86(10), 5697–5707. 10.1128/JVI.00146-12

63. Shukla, P., Nguyen, H. T., Torian, U., Engle, R. E., Faulk, K., Dalton, H. R., Bendall, R. P., Keane, F. E., Purcell, R. H., & Emerson, S. U. (2011). Cross-species infections of cultured cells by hepatitis E virus and discovery of an infectious virus-host recombinant. Proceedings of the National Academy of Sciences of the United States of America. 10.1073/pnas.1018878108

64. Silvin, A., Yu, C. I., Lahaye, X., Imperatore, F., Brault, J.-B., Cardinaud, S., Becker, C., Kwan, W.-H., Conrad, C., Maurin, M., Goudot, C., Marques-Ladeira, S., Wang, Y., Pascual, V., Anguiano, E., Albrecht, R. A., Iannacone, M., García-Sastre, A., Goud, B.,… Manel, N. (2017). Constitutive resistance to viral infection in human CD141 ^+^ dendritic cells. Science Immunology, 2(13), eaai8071. 10.1126/sciimmunol.aai8071

65. Songtanin, B., Molehin, A. J., Brittan, K., Manatsathit, W., & Nugent, K. (2023). Hepatitis E Virus Infections: Epidemiology, Genetic Diversity, and Clinical Considerations. Viruses, 15(6), 1389. 10.3390/v15061389

66. Sridhar, S., Yip, C. C. Y., Wu, S., Cai, J., Zhang, A. J.-X., Leung, K.-H., Chung, T. W. H., Chan, J. F. W., Chan, W.-M., Teng, J. L. L., Au-Yeung, R. K. H., Cheng, V. C. C., Chen, H., Lau, S. K. P., Woo, P. C. Y., Xia, N.-S., Lo, C.-M., & Yuen, K.-Y. (2018). Rat Hepatitis E Virus as Cause of Persistent Hepatitis after Liver Transplant. Emerging Infectious Diseases, 24(12), 2241–2250. 10.3201/eid2412.180937

67. Sumpter, R., Loo, Y.-M., Foy, E., Li, K., Yoneyama, M., Fujita, T., Lemon, S. M., & Gale, M. (2005). Regulating Intracellular Antiviral Defense and Permissiveness to Hepatitis C Virus RNA Replication through a Cellular RNA Helicase, RIG-I. Journal of Virology, 79(5), 2689–2699. 10.1128/JVI.79.5.2689-2699.2005

68. Swiecki, M., Gilfillan, S., Vermi, W., Wang, Y., & Colonna, M. (2010). Plasmacytoid Dendritic Cell Ablation Impacts Early Interferon Responses and Antiviral NK and CD8+ T Cell Accrual. Immunity. 10.1016/j.immuni.2010.11.020

69. Tam, A. W., Smith, M. M., Guerra, M. E., Huang, C.-C., Bradley, D. W., Fry, K. E., & Reyes, G. R. (1991). Hepatitis E virus (HEV): Molecular cloning and sequencing of the full-length viral genome. Virology, 185(1), 120–131. 10.1016/0042-6822(91)90760-9

70. Todt, D., François, C., Anggakusuma, Behrendt, P., Engelmann, M., Knegendorf, L., Vieyres, G., Wedemeyer, H., Hartmann, R., Pietschmann, T., Duverlie, G., & Steinmann, E. (2016). Antiviral Activities of Different Interferon Types and Subtypes against Hepatitis E Virus Replication. Antimicrobial Agents and Chemotherapy, 60(4), 2132–2139. 10.1128/AAC.02427-15

71. Venet, M., Ribeiro, M. S., Décembre, E., Bellomo, A., Joshi, G., Nuovo, C., Villard, M., Cluet, D., Perret, M., Pescamona, R., Paidassi, H., Walzer, T., Allatif, O., Belot, A., Trouillet-Assant, S., Ricci, E. P., & Dreux, M. (2023). Severe COVID-19 patients have impaired plasmacytoid dendritic cell-mediated control of SARS-CoV-2. Nature Communications, 14(1), 694. 10.1038/s41467-023-36140-9

72. Wang, W., Xu, L., Brandsma, J. H., Wang, Y., Hakim, M. S., Zhou, X., Yin, Y., Fuhler, G. M., Van Der Laan, L. J. W., Van Der Woude, C. J., Sprengers, D., Metselaar, H. J., Smits, R., Poot, R. A., Peppelenbosch, M. P., & Pan, Q. (2016). Convergent Transcription of Interferon-stimulated Genes by TNF-α and IFN-α Augments Antiviral Activity against HCV and HEV. Scientific Reports, 6(1), 25482. 10.1038/srep25482

73. Webster, B., Assil, S., & Dreux, M. (2016). Cell-Cell Sensing of Viral Infection by Plasmacytoid Dendritic Cells. Journal of Virology. 10.1128/jvi.01692-16

74. Webster, B., Werneke, S. W., Zafirova, B., This, S., Coléon, S., Décembre, E., Paidassi, H., Bouvier, I., Joubert, P.-E., Duffy, D., Walzer, T., Albert, M. L., & Dreux, M. (2018). Plasmacytoid dendritic cells control dengue and Chikungunya virus infections via IRF7-regulated interferon responses. eLife, 7, e34273. 10.7554/eLife.34273

75. Willemsen, J., Wicht, O., Wolanski, J. C., Baur, N., Bastian, S., Haas, D. A., Matula, P., Knapp, B., Meyniel-Schicklin, L., Wang, C., Bartenschlager, R., Lohmann, V., Rohr, K., Erfle, H., Kaderali, L., Marcotrigiano, J., Pichlmair, A., & Binder, M. (2017). Phosphorylation-Dependent Feedback Inhibition of RIG-I by DAPK1 Identified by Kinome-wide siRNA Screening. Molecular Cell, 65(3), 403–415.e8. 10.1016/j.molcel.2016.12.021

76. Wu, X., Dao Thi, V. L., Liu, P., Takacs, C. N., Xiang, K., Andrus, L., Gouttenoire, J., Moradpour, D., & Rice, C. M. (2018). Pan-Genotype Hepatitis E Virus Replication in Stem Cell–Derived Hepatocellular Systems. Gastroenterology, 154(3), 663-674.e7. 10.1053/j.gastro.2017.10.041

77. Yarilina, A., Park-Min, K.-H., Antoniv, T., Hu, X., & Ivashkiv, L. B. (2008). TNF activates an IRF1-dependent autocrine loop leading to sustained expression of chemokines and STAT1-dependent type I interferon–response genes. Nature Immunology, 9(4), 378–387. 10.1038/ni1576

78. Yin, X., Li, X., Ambardekar, C., Hu, Z., Lhomme, S., & Feng, Z. (2017). Hepatitis E virus persists in the presence of a type III interferon response. PLOS Pathogens, 13(5), e1006417. 10.1371/journal.ppat.1006417

79. Yin, X., Ying, D., Lhomme, S., Tang, Z., Walker, C. M., Xia, N., Zheng, Z., & Feng, Z. (2018). Origin, antigenicity, and function of a secreted form of ORF2 in hepatitis E virus infection. Proceedings of the National Academy of Sciences of the United States of America. 10.1073/pnas.1721345115

80. Yu, C., Boon, D., McDonald, S. L., Myers, T. G., Tomioka, K., Nguyen, H., Engle, R. E., Govindarajan, S., Emerson, S. U., & Purcell, R. H. (2010). Pathogenesis of Hepatitis E Virus and Hepatitis C Virus in Chimpanzees: Similarities and Differences. Journal of Virology, 84(21), 11264–11278. 10.1128/JVI.01205-10

81. Yun, T. J., Igarashi, S., Zhao, H., Perez, O. A., Pereira, M. R., Zorn, E., Shen, Y., Goodrum, F., Rahman, A., Sims, P. A., Farber, D. L., & Reizis, B. (2021). Human plasmacytoid dendritic cells mount a distinct antiviral response to virus-infected cells. Science Immunology, 6(58), eabc7302. 10.1126/sciimmunol.abc7302

82. Zhong, P., Agosto, L. M., Ilinskaya, A., Dorjbal, B., Truong, R., Derse, D., Uchil, P. D., Heidecker, G., & Mothes, W. (2013). Cell-to-Cell Transmission Can Overcome Multiple Donor and Target Cell Barriers Imposed on Cell-Free HIV. PLoS ONE, 8(1), e53138. 10.1371/journal.pone.0053138

83. Zhou, X., Xu, L., Wang, W., Watashi, K., Wang, Y., Sprengers, D., De Ruiter, P. E., Van Der Laan, L. J. W., Metselaar, H. J., Kamar, N., Peppelenbosch, M. P., & Pan, Q. (2016). Disparity of basal and therapeutically activated interferon signalling in constraining hepatitis E virus infection. Journal of Viral Hepatitis, 23(4), 294–304. 10.1111/jvh.12491

