## Supplementary figure for "Hepatitis E Virus-induced antiviral response by plasmacytoid dendritic cells is modulated by the ORF2 protein"

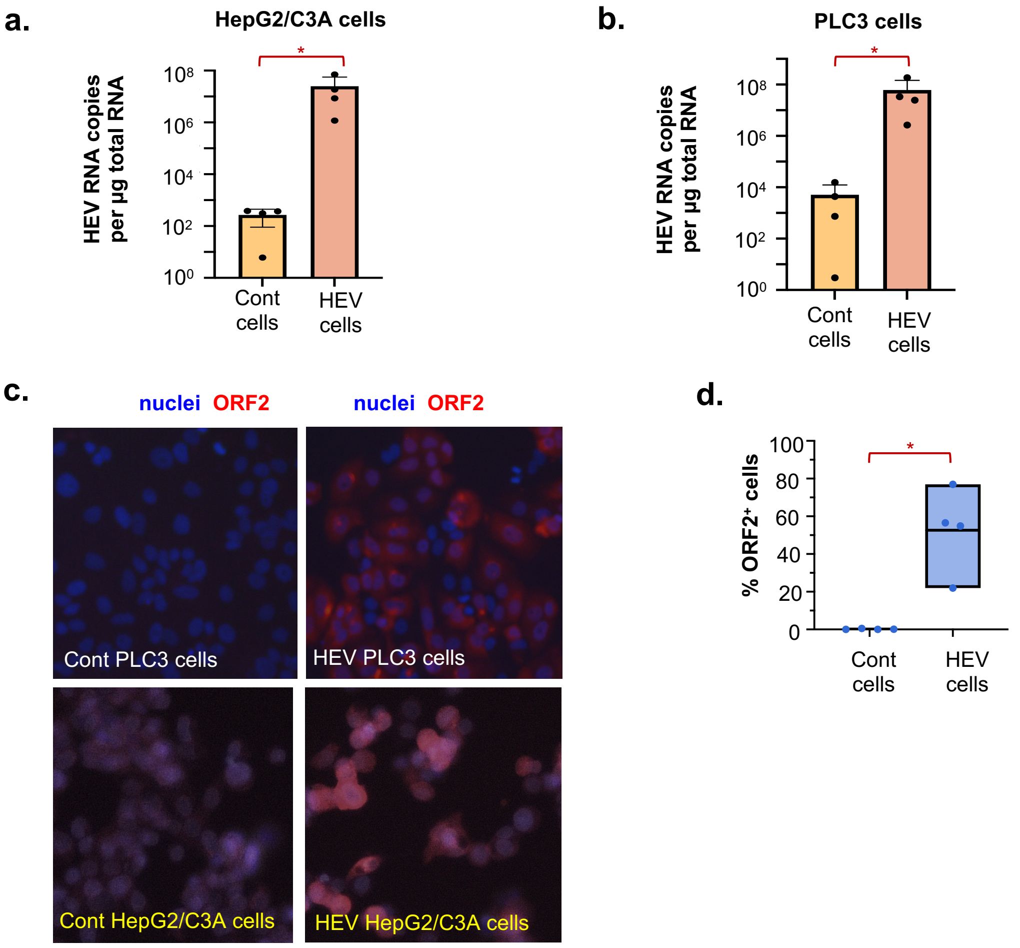


e.

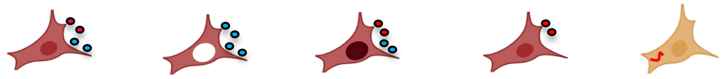

| WT/Mutant | | WT cells | 5R/5A cells | NES 12 cells | STOP cells | GAD cells |
|---|---|---|---|---|---|---|
| Extracellular | Virions (ORF2i) | ✓ | ✕ | ✓ (-) | ✓ | ✕ |
| | RNA | ✓ | Non encapsidated (non-infectious) | ✓ | ✓ | ✕ |
| | ORF2g/c | ✓ | ✓(+++) | ✓ | ✕ | ✕ |
| Intracellular | Virions (ORF2i) | ✓ | ✕ | ✓ (-) | ✓ | ✕ |
| | RNA | ✓ | ✓ | ✓ | ✓ | ✓ (non-replicating) |
| | ORF2i | ✓ | ✕ | ✓ | ✓ | ✕ |
| | Nuclear ORF2 | ✓ | ✕ | ✓(+++) | ✓ | ✕ |

**Supplementary Figure 1**

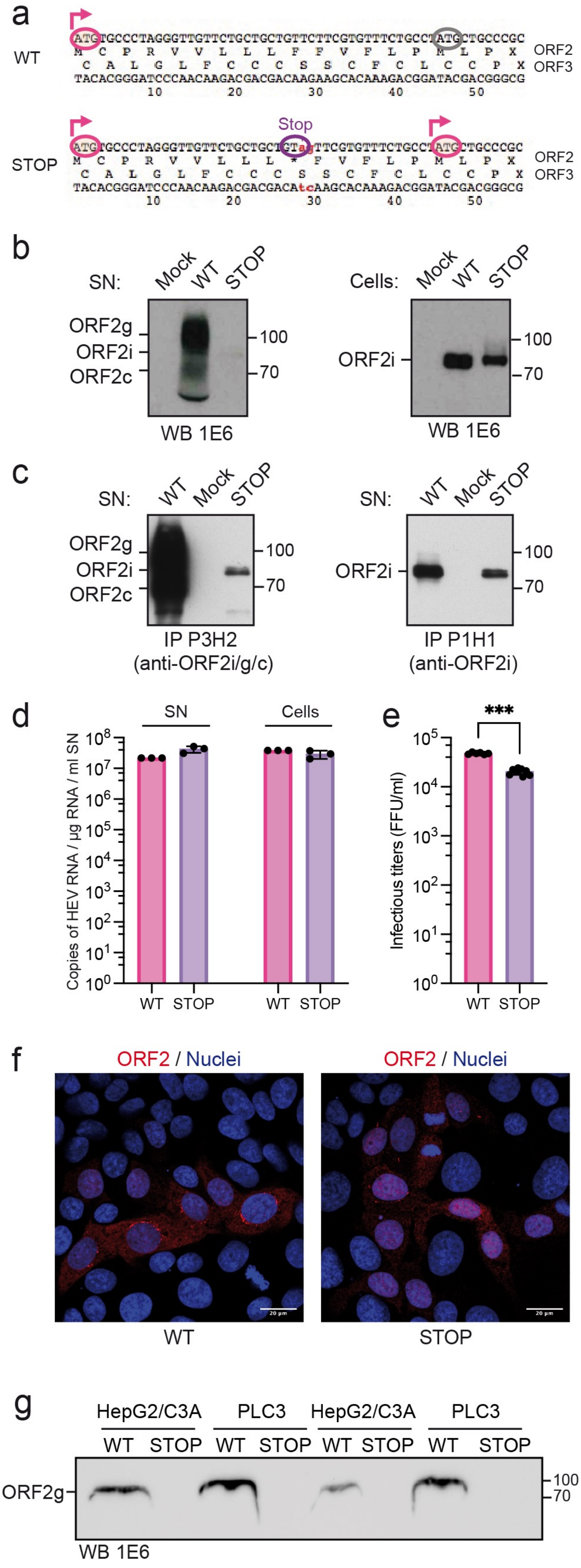


**Supplementary Figure 2**

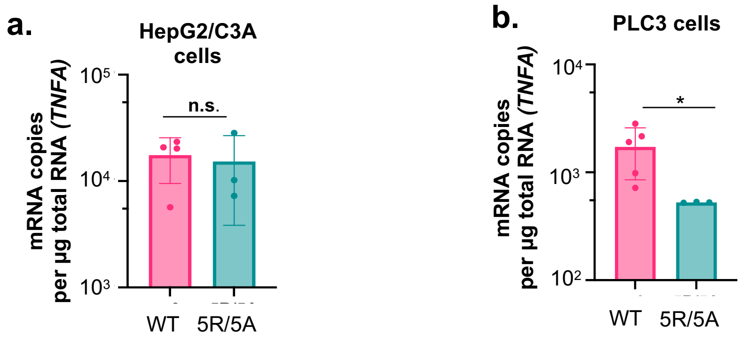


Supplementary Figure 3

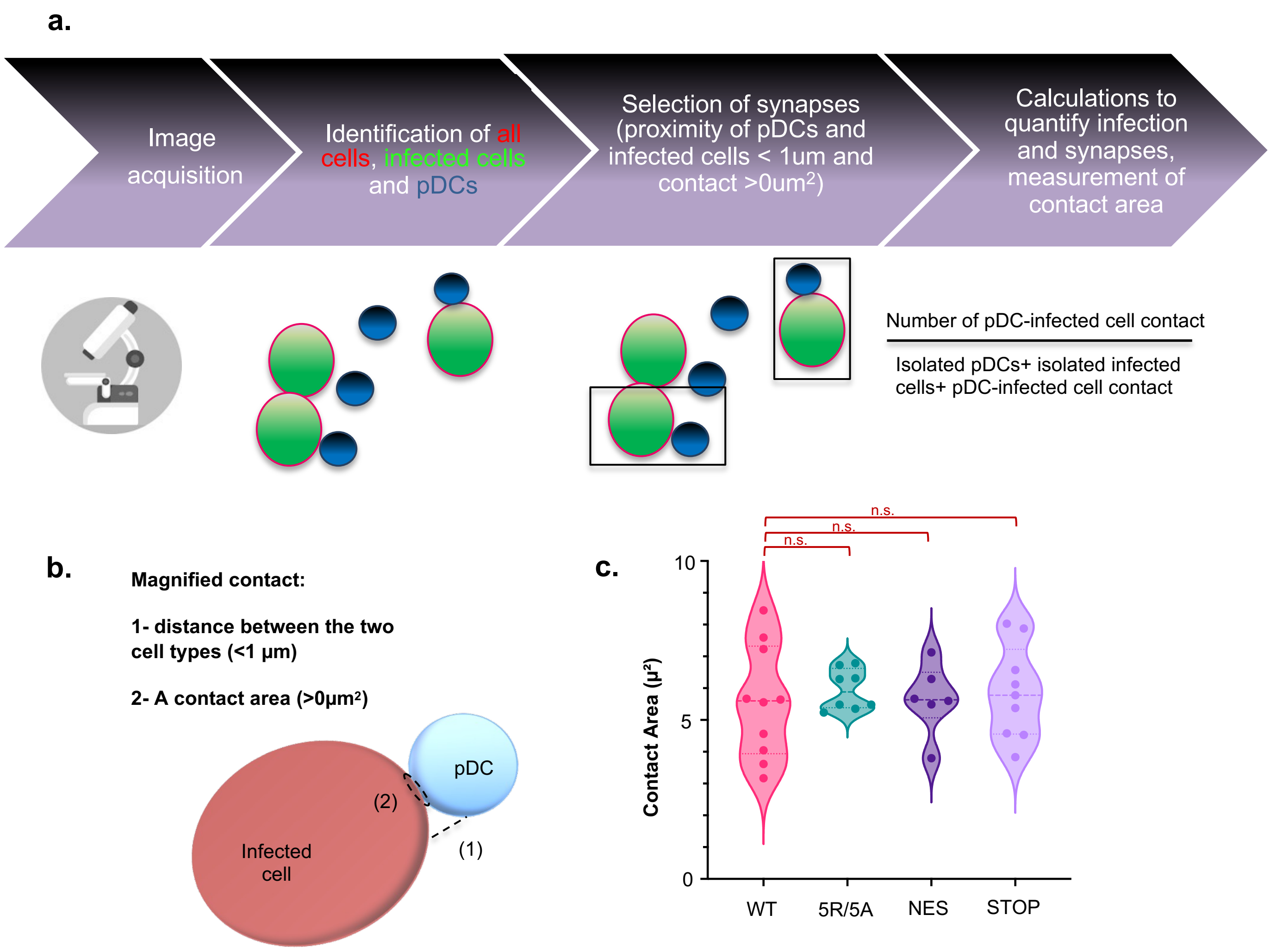


**Supplementary Figure 4**
