## Supplementary legend for "Hepatitis E Virus-induced antiviral response by plasmacytoid dendritic cells is modulated by the ORF2 protein"

**Legend: Supplementary figures**

**Supplementary Figure 1.** (**A-B**) Quantification of the replication levels of HEV at 6 days post electroporation with 10µg RNA [HEV cells] or mock electroporation without RNA [Cont cells] (as described in Material and method section) by RT-qPCR detection of HEV RNA (amplicon/primers designed in ORF2) in HepG2/C3A **(A)** and PLC3 cells **(B)**, respectively; means ± SD; each dot represents one independent experiment; n=4 for PLC3 cells and HepG2/C3A cells; statistical analysis was done using Wilcoxon rank sum test with continuity correction and p value adjustment with Bonferroni method; *p* values as: * ≤0.05, ** ≤0.005 and *** ≤0.0005. **(C)** HEV infection levels determined by immunodetection of ORF2 protein using 1E6 antibody, are presented as representative images in HEV-producing PLC3 and HepG2/C3A cells *versus* control cells [cont] transfected with no RNA as negative control, at 6 d.p.e. **(D)** Quantification by flow cytometry of the frequency of HEV ORF2-positive PLC3 cells. Statistical analysis was done using Wilcoxon rank sum test with continuity correction and p value adjustment with FDR method; *p* values as: * ≤0.05, ** ≤0.005 and *** ≤0.0005 **(E)** Illustrative table describing the differences between WT and mutants in terms of intracellular and extracellular viral products in electroporated cells

**Supplementary Figure 2. Determinant of the production of** **ORF2g/c forms.** **(A)** Construction: a stop codon introduced in the ORF2 signal peptide without affecting ORF3 expression. **(B)** WB detection of ORF2 in the supernatant [SN] and cell lysates [cells] of HEV-producing PLC3 cells (using 1E6 mAb: detect all ORF2 species, mostly ORF2g/c in WT SN). **(C)** Immunoprecipitation in SN of all ORF2 forms *versus* ORF2i using P3H2 and P1H1 mAbs, respectively. (**D-E).** Quantification of HEV RNA levels in SN and HEV-infected PLC3 cells and extracellular infectious titers; mean±SD; *n*≥*3.* **(F)** Imaging of ORF2 (1E6 mAb) in Huh-7.5 cells infected by WT and STOP mutant. **(G)** Western blot detection of ORF2 in the SNs of HEV-replicating HepG2/C3A and PLC3 cells 6d.p.e. with HEV WT or STOP RNA in two independent experiments (using 1E6 mAb, which detects all ORF2 species, mostly ORF2g/c in WT SN).

**Supplementary Figure 3. Differences in HEV-induced immune signaling among cell types.** **(A-B)** Quantification of the transcript expression levels of *TNF* determined at 6 d.p.e. of HepG2/C3A (**A**) and PLC3 **(B)** cells. Bars represent copy number per µg total RNA; means ± SD; each dot represents one independent experiment (n=3 to 5). Statistical analysis was done using Wilcoxon rank sum test with continuity correction; *p* values as: * ≤0.05, ** ≤0.005 and *** ≤0.0005. *OAS2* mRNA abundance in HEV-replicating cells, in the presence or absence of pDCs.

**Supplementary Figure 4. Confocal imaging pipeline to quantify cell proximity between pDCs and infected cells. (A)** Illustrative pipeline explaining steps in acquisition and treatment of confocal images used to quantify infected cells, enumerate the contacts between infected cells and pDCs, and measure the contact area. **(B)** Illustration describing criteria of contact enumeration as set in the macro used to analyze the confocal images. **(C)** Violin plots represent area of contact/proximity of PLC3 infected cells and pDCs from automatic tracking (*i.e.,* defined within a cell-to-cell distance < 1 µm; contact area> 0µm^2^) with detection of CTV/pDCs and CMTPX^+^ ORF2^+^/infected cells, as reference; violin plots present median and each dot for independent image (n = 3). Statistical analysis was done using Wilcoxon rank sum test.
