## Supplementary table for "Hepatitis E Virus-induced antiviral response by plasmacytoid dendritic cells is modulated by the ORF2 protein"

**Supplementary Table 1. Primer used for RT-qPCR**

| Name | Forward/Reverse | Specificity | Sequence 5’-3’ |
| --- | --- | --- | --- |
| HEV | Forward | HEV | GGT GGT TTC TGG GGT GAC |
|  | Reverse | HEV | AGG GGT TGG TTG GAT GAA |
| GAPDH | Forward | human | AGGTGAAGGTCGGAGTCAACG |
|  | Reverse | human | TGGAAGATGGTGATGGGATTTC |
| Interferon lambda 1 IL29 | Forward | human | TCCTAGACCAGCCCCTTCA |
|  | Reverse | human | GTGGGCTGAGGCTGGATA |
| ISG15 | Forward | human | GACAAATGCGACGAACCTCT |
|  | Reverse | human | CGGCCCTTGTTATTCCTCA |
| ISG56 | Forward | human | GGGCAGACTGGCAGAAG |
|  | Reverse | human | CTATAGCGGAAGGGATTTGA |
| MXA | Forward | human | ACAGGACCATCGGAATCTTG |
|  | Reverse | human | CCCTTCTTCAGGTGGAACAC |
| TNF alpha | Forward | human | AGATGATCTGACTGCCTGGG |
|  | Reverse | human | CTGCTGCACTTTGGAGTGAT |
| IL6 | Forward | human | GTCAGGGGTGGTTATTGCAC |
|  | Reverse | human | AGTGAGGAACAAGCCAGAGC |
| OAS2 | Forward | human | CCTGAAGCCCTACGAAGA |
|  | Reverse | human | TTAAGACTGTTTTCCGTCCA |
